# Induced Estrogen Receptor SUMOylation drives SERD activity

**DOI:** 10.64898/2026.06.26.734426

**Authors:** Matthias Hinterndorfer, Caroline Schätz, Stefan Schmitt, Martin Schönlein, David M. Hoi, Izabela Krecioch, Fabian Frommelt, Mykhailyna Shlei, Lukas Kater, Martin Pacesa, Miquel Munoz, Georg Kempf, Katharina Kladnik, Paul Batty, Hana Imrichova, Jacob D. Aguirre, Sandra Högler, Simone Cavadini, Davide Seruggia, Bruno E. Correia, Anna C. Obenauf, Nicolas H. Thomä, Georg E. Winter

**Affiliations:** AITHYRA Research Institute for Biomedical Artificial Intelligence of the Austrian Academy of Sciences, Vienna, Austria; CeMM Research Center for Molecular Medicine of the Austrian Academy of Sciences, Vienna, Austria; Friedrich Miescher Institute for Biomedical Research, Basel, Switzerland; Faculty of Science, University of Basel, Basel, Switzerland; Research Institute of Molecular Pathology (IMP), Vienna BioCenter (VBC), Vienna, Austria; Discovery Sciences, Novartis Biomedical Research, Basel, Switzerland; Institute for Cancer Research (ISREC), Swiss Federal Institute of Technology Lausanne (EPFL), Lausanne, Switzerland; Institute of Pharmacology and Toxicology, University of Zurich, Switzerland; St. Anna Children’s Cancer Research Institute (CCRI), Vienna, Austria; University of Veterinary Medicine Vienna, Center for Pathobiology, Vienna, Austria

## Abstract

The ligand dependent transcription factor estrogen receptor α (ERα) is a key driver of and important drug target in breast cancer. Patients with advanced disease are typically treated with selective ERα degraders (SERDs), whose therapeutic activity is commonly attributed to induced ERα protein degradation. Yet the exact mechanism and relevance of degradation for clinical efficacy remain unclear. We show that SERDs directly induce ERα SUMOylation, thereby triggering degradation via SUMO-targeted ubiquitin ligases (STUbLs). Inactivation of STUbLs prevents ERα degradation and counterintuitively further sensitizes breast cancer cells to SERDs, rather than conferring resistance. SERD efficacy is independent of ERα degradation, challenging the degradation-centric model of SERD action. Instead, SUMOylation recruits transcriptional co-repressors, turning SUMOylated ERα into a ‘dominant-negative’ repressor. Thus, SUMOylation rather than degradation is the direct consequence and driver of SERD activity. Our findings position SUMO-inducing drugs as a hitherto underappreciated yet clinically validated therapeutic modality with broad applicability.

## Main Text

ERα⁺ breast cancer is the most common breast cancer subtype in women and a leading cause of cancer-related mortality (*1, 2*). These tumors depend on ERα, a ligand-inducible nuclear hormone receptor that upon estrogen binding activates pro-proliferative transcriptional programs through recruitment of co-regulators and chromatin-modifying enzymes (*3–5*). Accordingly, therapeutic suppression of ERα signaling has been a central strategy in the treatment of hormone receptor–positive breast cancer for more than three decades.

ERα⁺ breast cancers are initially treated with aromatase inhibitors that reduce endogenous estrogen levels to limit ERα activation, or with selective estrogen receptor modulators (SERMs) such as Tamoxifen, which competitively bind to ERα and inhibit its transcriptional activity. Advanced, relapsing or resistant patients are typically switched to Fulvestrant or Elacestrant, which show more complete ERα antagonism and superior efficacy especially in patients with ERα mutations or endocrine resistance. In addition to ERα inhibition, these compounds also induce the proteasomal degradation of ERα protein, establishing the class of selective ERα degraders (SERDs).

However, despite over two decades of widespread clinical use and substantial investment into the development of next-generation SERDs with improved pharmacokinetic properties, the mechanism of action (MoA) of SERDs remains incompletely understood. In particular, as the cellular machinery inducing ERα degradation has largely remained elusive, we lack a definitive understanding of the relevance of ERα degradation for the clinical efficacy of SERDs (*6*). Correlative observations suggest that ERα degradation and cytostatic anticancer activity might be disconnected (*7–13*). However, in the absence of definitive, causal evidence, the prevailing view in the field remains that SERDs exert their effects largely through induced ERα degradation (*14–17*).

To address these critical mechanistic gaps for SERDs, we systematically mapped effectors of SERD-induced ERα degradation and SERD efficacy using orthogonal functional genomics and proteomics screens combined with biochemical reconstitutions. SERD treatment has previously been linked to ERα SUMOylation in breast cancer cells (*7, 9, 12, 18*). Here, we find that SERDs functionalize a native association of ERα with PIAS SUMO E3 ligases to directly promote SUMOylation. SUMOylation, in turn, triggers the activity of SUMO-targeting ubiquitin ligases (STUbLs), resulting in proteasomal degradation. Inactivation of ERα degradation by genetic STUbL depletion does not protect breast cancer cells from SERD-induced growth arrest but instead enhances SERD efficacy. This, for the first time proves that the antiestrogenic and cytostatic activity of SERDs is completely independent of ERα degradation. Mechanistically, we show that ERα SUMOylation assembles a repressive transcriptional environment that actively silences estrogen signaling and disrupts breast cancer cell growth more effectively than ERα protein degradation. SUMOylated ERα thus drives transcriptional repression in a ‘dominant-negative’ fashion. Our findings position induced SUMOylation as conceptually novel yet clinically already established therapeutic modality for the silencing of oncogenic transcriptional programs.

### SERDs induce ERα degradation via SUMO-targeted ubiquitin E3 ligases

To map the molecular machinery involved in SERD activity, we set up two complementary flow cytometry-based CRISPR screens for effectors of ERα degradation: a focused screen targeting members of the ubiquitin-proteasome system (UPS) in the ERα positive breast adenocarcinoma cell line MCF7 and a genome-wide screen in KBM7, a well-established model for functional genetic screens (*19, 20*). We expressed a dual-fluorescence ERα protein stability reporter, consisting of ERα-GFP or ERα-BFP fusions coupled to mCherry for normalization (Table S1), in cell lines carrying doxycycline (dox)-inducible Cas9 alleles. We mutagenized these reporter cells with sgRNA libraries, treated them with SERDs or control compounds, and isolated cells with elevated or decreased ERα levels via fluorescence activated cell sorting (FACS; Fig. 1A, Table S2). Screens in both cell lines yielded similar and partially overlapping top hits. Screens performed in the absence of SERDs identified the E3 ligase UBR5 and its regulator OTUD5 (*21*) (fig. S1A and B), recapitulating the native degradation pathway triggered by ERα agonists (*22, 23*). UBR5 and OTUD5 similarly scored in a screen performed with AZD9496, a SERD known to enhance the endogenous ERα degradation pathway (*22*) (fig. S1C). In comparison, screens performed after treatment with Fulvestrant (Ful) or the clinical phase III investigational SERDs Camizestrant (Cami) or Giredestrant (Gire) (*24, 25*) (Fig. 1B) identified a partially overlapping group of enzymes involved in the conjugation and sensing of the small ubiquitin-like modifier SUMO (Fig. 1C, fig. S1D-I).

**Fig. 1.**
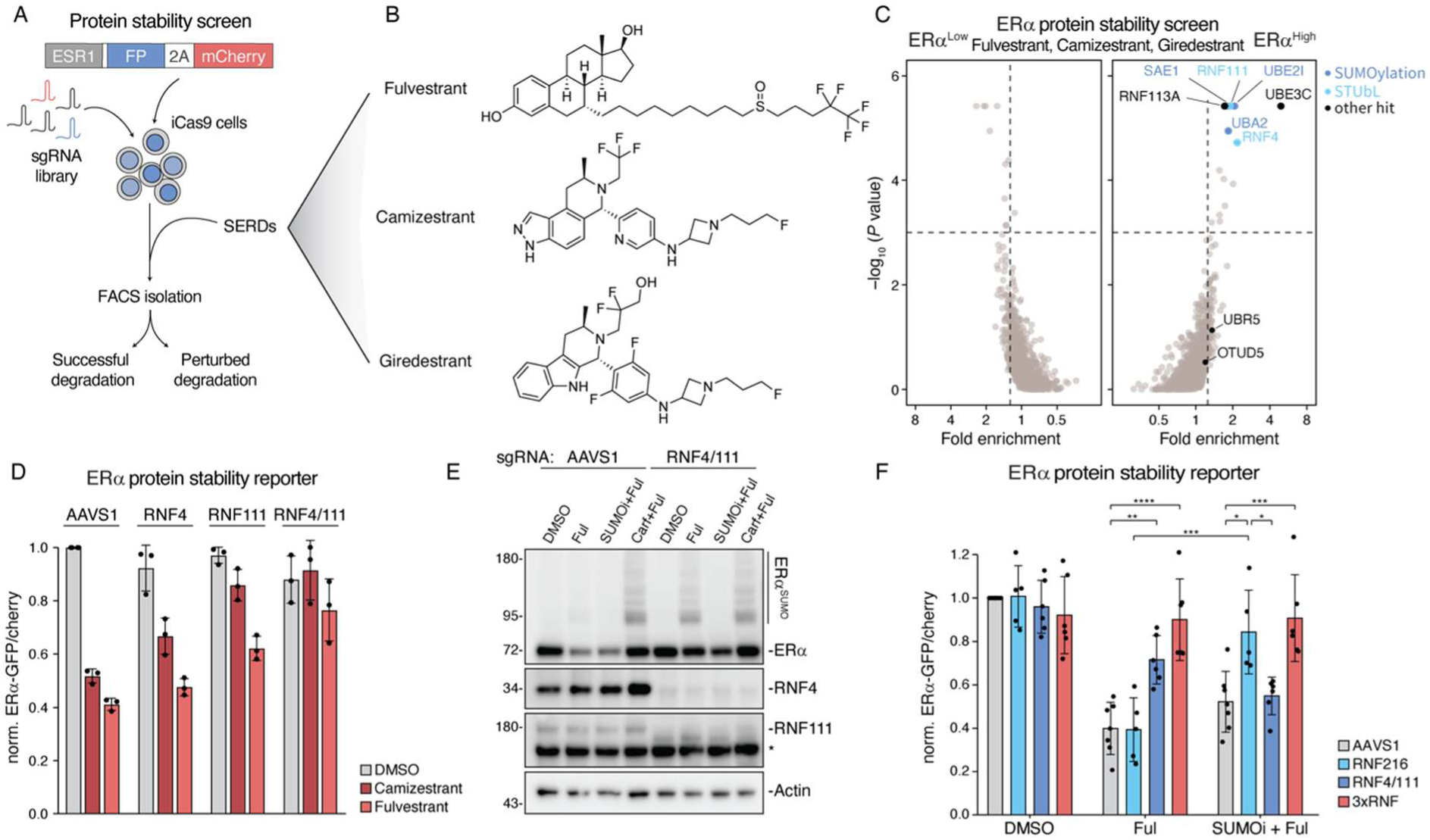
SERDs induce ERα degradation via SUMO-targeted E3 ligases. **(A)** Schematic of FACS-based ERα protein stability CRISPR-Cas9 screens. Doxycycline (Dox)-inducible Cas9 (iCas9) MCF7 ERα-GFP or KBM7 ERα-BFP reporter cells were transduced with ubiquitin-proteasome system (UPS)-focused or genome-wide sgRNA libraries, respectively, treated with SERDs and sorted based on ERα/mCherry ratios. **(B)** Structures of the SERDs Fulvestrant, Camizestrant, Giredestrant used in ERα stability CRISPR screens. **(C)** FACS-based CRISPR screens for ERα protein stability. MCF7 iCas9 ERα-GFP reporter cells were treated with DMSO or indicated SERDs (10 nM each) for 6 h before sorting. Gene-level fold enrichment and one-sided MAGeCK P value were calculated across screens performed with Fulvestrant, Camizestrant and Giredestrant; n = 2 biological replicates each. SUMO gene variants and SUMOylation enzymes are highlighted in blue, SUMO-targeted ubiquitin E3 ligases (STUbLs) in cyan. Screen results and enrichment scores for individual screens in fig S1A-B and fig. S1D-F. **(D)** FACS-based screen validation. MCF7 iCas9 ERα-GFP reporter cells were transduced with AAVS1-, RNF4- or RNF111-targeting sgRNAs either individually or simultaneously, treated with DMSO, Fulvestrant or Camizestrant (both 10 nM) for 6 h, and ERα-GFP signal normalized by mCherry was quantified by FACS. n = 3 independent experiments, mean ± sd. **(E)** Immunoblot-based screen validation. MCF7 iCas9 cells were transduced with AAVS1-or co-transduced with RNF4- and RNF111-targeting sgRNAs, treated with DMSO or Fulvestrant (10 nM) for 6 h with or without 1 h pre-treatment with ML792 (SUMOi; 2 μM) or Carfilzomib (Carf; 1 μM). Protein levels were analysed via immunoblot. Data representative of n = 3 independent experiments. **(F)** RNF216-based degradation. MCF7 iCas9 ERα-GFP reporter cells were transduced with AAVS1-, RNF216-, combined RNF4- and RNF111- or combined RNF4-, RNF111- and RNF216-targeting (3xRNF) sgRNAs, pre-treated for 1 h with or without ML792 (SUMOi; 2 μM) followed by treatment with DMSO or Fulvestrant (10 nM) for 6 h. ERα-GFP signal was quantified by FACS as in d. n = 7 (AAVS1), 6 (RNF4/111, 3xRNF) or 5 (RNF216) independent experiments, mean ± sd. Two-way ANOVA with Tukey HSD post-hoc test, *p<0.05, **p<0.01, ***p<0.001, ****p<0.0001 *ESR1*, Estrogen Receptor 1; FP, (Green or Blue) Fluorescent Protein; Ful, Fulvestrant; Carf, Carfilzomib; SUMOi, ML792.

SUMO is a small, ubiquitin-like protein that is post-translationally attached to a wide range of target proteins, including ERα (*26*), regulating many cellular functions including DNA damage response, transcription and proteostasis (*27–29*). Like ubiquitylation, SUMOylation is conducted by a cascade of a dedicated SUMO E1 activating enzyme, a SUMO E2 conjugating enzyme, and several SUMO E3 ligases. Beyond its diverse regulatory roles, SUMOylation can also promote proteasomal turnover via the activity of dedicated SUMO-targeted ubiquitin E3 ligases (STUbLs), which recognize SUMOylated proteins and mark them for degradation (*30, 31*). In screens in MCF7 cells, we identified the core SUMOylation machinery, including the heterodimeric SUMO E1 enzyme SAE1/UBA2, the SUMO E2 UBE2I (UBC9) and the two canonical STUbLs RNF4 and RNF111 as modulators of SERD-induced ERα degradation (*32–34*). Genome-wide screens in KBM7 cells yielded RNF4 in addition to the recently discovered STUbL TOPORS and another ubiquitin E3 ligase, RNF216. Despite their consistent score in the CRISPR screens, perturbation of individual STUbLs had only minor effects on SERD-induced ERα degradation in MCF7 cells (fig. S2A and B). We thus reasoned that their role in ERα degradation might be functionally redundant. Combined knockout (KO) of RNF4 and RNF111 indeed displayed additive effects, preventing degradation of the ERα stability reporter as well as endogenous ERα (Fig. 1D and E). In contrast to combined KO of RNF4 and RNF111, KO of the non-redundant SUMO E1 (SAE1/UBA2) or E2 (UBE2I) enzymes or pre-treatment with the SUMOylation inhibitor ML792 (SUMOi) barely influenced ERα degradation (Fig. 1E and fig. S2A-C), even though SUMOylation was efficiently blocked (Fig. 1E, fig. S2C). We thus speculated that a compensatory pathway might maintain ERα degradation in the absence of SUMOylation. To this end, we tested the impact of other screen hits on ERα degradation after SUMOi treatment. While RNF216 KO did not influence ERα degradation by Fulvestrant in unperturbed MCF7 cells, it mediated ERα degradation in the absence of functional SUMOylation networks (Fig. 1F and fig. S2D). This indicated a switch from primarily STUbL-dependent degradation in unperturbed cells to an RNF216-dependent backup pathway activated in the absence of SUMOylation. Combined KO or knockdown of RNF4, RNF111 and RNF216 (3xRNF) invariably prevented ERα degradation in MCF7 as well as T47D cells independent of the presence or absence of SUMOylation (Fig. 1F and fig. S2D-F). Together, these data show that SERDs drive ERα degradation primarily via the redundant action of the STUbLs RNF4 and RNF111, with RNF216 providing a failsafe mechanism triggered in the absence of SUMOylation.

### PIAS family ligases SUMOylate ERα

In agreement with previous observations reporting cellular ERα SUMOylation upon SERD treatment (*7, 9, 12, 18*), we observed the appearance of high molecular weight species on ERα immunoblots indicative of SERD-induced SUMOylation (Fig. 1E and fig. S2C-F). Using ERα immunoprecipitation as well as a split-HaloTag-based (*35*) live cell SUMOylation assay, we confirmed ERα SUMOylation upon SERD but not SERM (4-Hydroxytamoxifen, 4-OHT) treatment (Fig. 2A and fig. S3A and B). ERα SUMOylation occurred in various breast cancer cell lines (fig. S3C), set on rapidly after SERD treatment, temporally preceded ERα degradation (fig. S3D), and was stable only after treatment with the proteasome inhibitor Carfilzomib or the ubiquitylation inhibitor TAK243 (Fig. 1E, fig. S2D and fig. S3C and D). Combined RNF4 and RNF111 KO stabilized SUMOylated ERα similar to proteasome inhibition (Fig. 1E and fig. S2D-F), together highlighting the rapid ubiquitylation and turnover of SUMOylated ERα by the STUbLs RNF4/RNF111. We thus next developed fully recombinant *in vitro* ubiquitylation assays to test whether STUbL-based ubiquitylation is directly triggered by SERD binding or is a consequence of induced ERα-SUMOylation. Incubating RNF4 or RNF111 together with the core ubiquitylation machinery, we observed robust ubiquitylation of ERα N-terminally fused to two SUMO2 moieties (2×SUMO2-ERα) irrespective of the presence of Fulvestrant or Estradiol. In contrast, no ubiquitylation activity was observed toward non-SUMOylated ERα (maltose binding protein (MBP)-ERα; Fig. 2B). This suggested that STUbL-mediated ERα ubiquitylation is independent of the bound ligand and instead solely depends on prior SUMOylation (*36, 37*).

**Fig. 2.**
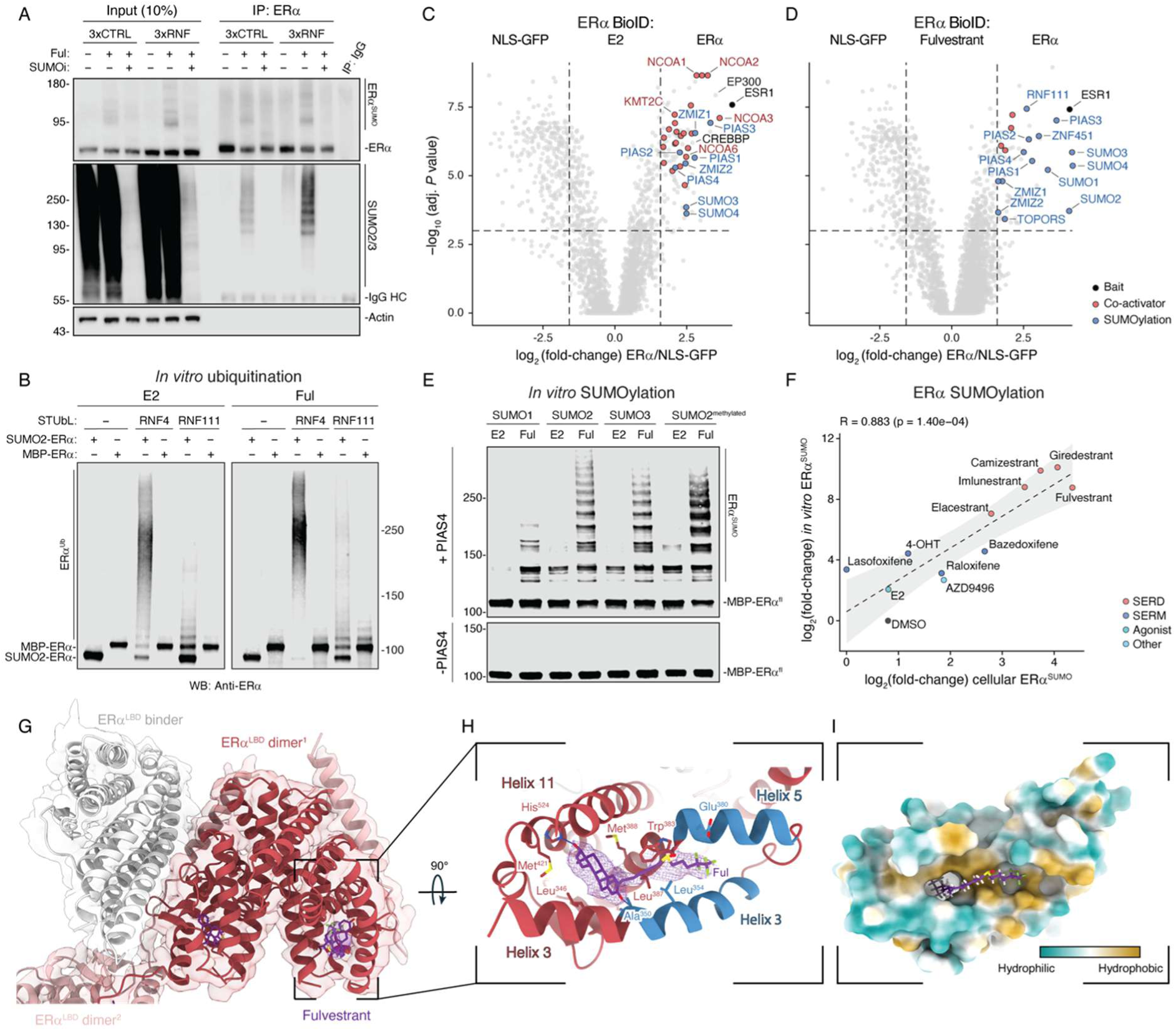
SERDs trigger ERα SUMOylation by the PIAS SUMO E3 ligase family. **(A)** ERα immunoprecipitation. MCF7 cells were pre-treated with Carfilzomib (1 μM) for 1 hour, followed by treatment with DMSO, Fulvestrant (10 nM) or Fulvestrant plus SUMOi (2 μM) for 6 hours. ERα was immunopurified using an ERα-specific antibody, followed by immunoblotting with a SUMO2/3-specific antibody. Data representative of n = 3 independent experiments. **(B)** *In vitro* ubiquitylation assay. *In vitro* ubiquitylation of full-length ERα, purified either as an MBP fusion or fused N-terminally to two SUMO2 moieties, was performed in the presence of Estradiol or Fulvestrant and either RNF4 or RNF111. Ubiquitylated ERα was visualized via ERα immunoblotting. Data representative of n = 2 independent experiments. **(C and D)** ERα BioID proximity labelling. ERα-miniTurbo cells were treated with 10 nM Fulvestrant (C) or Estradiol (D). Transcriptional activators (red; GOBP:0045944) and SUMO pathway components (blue; GOBP:0016925) in the scoring window (FC > 3, adj. P value < 0.001, one-way analysis of variance (ANOVA) with Tukey’s post hoc test and Benjamini-Hochberg correction, n = 3 biological replicates) are highlighted. **(E)** *In vitro* SUMOylation assay. Recombinantly purified full-length MBP-ERα, bound to Estradiol or Fulvestrant was incubated with the indicated SUMO isoforms and the core SUMOylation machinery in the presence or absence of PIAS4. SUMOylated ERα was visualized via ERα immunoblotting. Data representative of n = 2 independent experiments. **(F)** Correlation of cellular and *in vitro* ERα SUMOylation. ERα SUMOylation obtained by co-treatment of MCF7 cells with Carfilzomib (1 μM) and 10 nM ERα ligands was quantified via immunoblotting (fig. S3M) and correlated with *in vitro* SUMOylation of MBP-ERα^DBD-LBD^ complexed with indicated ligands (fig. S3N). Dashed line and ribbon, linear regression with 95% confidence interval. Pearson R, P value (Pearson’s two-sided correlation test) are shown. **(G)** Composite cryo-EM density map of the main ERα^LBD^ dimer^1^ (red) bound to Fulvestrant (purple) and a *de novo* designed ERα^LBD^ binder (gray), overlaid with the corresponding atomic model. **(H)** Close-up view of region outlined in (G). The steroid core is rotated by 180° compared to Estradiol (PDB ID: 1ERE) and forms a network of hydrogen bonds and hydrophobic interactions within the ligand-binding pocket. The Fulvestrant tail protrudes into the hydrophobic co-activator binding groove (blue). Cryo-EM density for Fulvestrant is shown as purple mesh. Protein side chains interacting with Fulvestrant (d < 4 Å) are shown, with hydrogen bonds represented by dotted lines. **(I)** Surface representation of monomeric ERα^LBD^. Colors represent molecular lipophilicity potential (MLP). The tail occupies the shallow hydrophobic co-activator binding groove, resulting in displacement of H12 from the LBD. E2, Estradiol; Ful, Fulvestrant; Cami, Camizestrant; MBP, maltose binding protein; STUbL, SUMO-targeted ubiquitin ligase; 4-OHT, 4-Hydroxytamoxifen; fl, ERα full-length protein.

We thus hypothesized that SERDs might act as direct inducers of ERα SUMOylation. While our screens readily identified the non-redundant SUMO E1 and E2 enzymes, they did not yield any SUMO E3 ligases, probably due to partial redundancy (*38, 39*). We turned to proximity-labelling to identify putative SUMO E3 ligases that might be recruited to ERα upon SERD binding. We expressed either GFP tagged with a nuclear localization signal (NLS-GFP) or ERα fused to a ‘miniTurbo’ biotin ligase (*40*) in MCF7 cells and, after a short pulse of biotin treatment, isolated biotinylated proteins for mass spectrometry-based proteomics. Compared to NLS-GFP, we observed the expected enrichment of transcriptional co-activators with ERα after Estradiol treatment, which were deprived in Fulvestrant-treated ERα (Fig. 2C and D, fig. S3E-G), consistent with abolished recruitment of the nuclear receptor co-activator NCOA1 observed in fluorescence polarization (FP) assays (fig. S3H). We furthermore found ERα associated with several SUMO E3 ligases, including all four members of the PIAS family (PIAS1-4), the PIAS-like ligases ZMIZ1-2 (*41, 42*) and ZNF451 (Fig. 2C and D and fig. S3E and G), indicating redundant involvement of multiple SUMO E3 ligases. Consequently, only simultaneous KO of multiple SUMO E3 ligases reduced the amount of ERα SUMOylation induced by Fulvestrant (fig. S3I). While SUMO protein variants (SUMO1-4), STUbLs and the SUMO protease SENP6 were enriched exclusively after Fulvestrant treatment, PIAS and PIAS-like ligases were also associated with ERα after Estradiol (E2) treatment (fig. S3G), consistent with their reported role in regulating endogenous ERα transcriptional activity (*7, 26*). Using cellular NanoBiT complementation (fig. S3J) as well as *in vitro* ERα pulldowns (fig. S3K), we confirmed the existence of an endogenous ERα:PIAS interaction, which was enhanced by Fulvestrant. Thus, SERDs induce SUMOylation by enhancing the native association of ERα with SUMO E3 ligases rather than by inducing *de novo* protein-protein interactions.

To test whether ERα SUMOylation is a direct biochemical consequence of SERD binding, we performed *in vitro* SUMOylation assays using recombinant proteins. We incubated MBP-ERα with the core SUMOylation machinery with or without PIAS SUMO E3 ligases in the presence of either Fulvestrant or Estradiol. While the core SUMOylation machinery alone was inactive, the addition of PIAS1 or PIAS4 induced low levels of SUMOylation of recombinant ERα complexed with Estradiol (Fig. 2E and fig. S3L). The SUMOylation activity of PIAS enzymes was drastically enhanced in the presence of Fulvestrant, displaying pronounced ERα decoration with SUMO1, -2 and -3 isoforms (Fig. 2E and fig. S3L) and recapitulating ERα SUMOylation patterns observed after SERD treatment in cells (Fig. 2E and fig. S3C). SUMOylation was unchanged with methylated SUMO2 incapable of chain formation, indicating multi-mono- rather than poly-SUMOylation of ERα (Fig. 2E and fig. S3L). Over a panel of ERα ligands, *in vitro* SUMOylation tightly correlated with cellular SUMOylation (Fig. 2F and fig. S3M and N), indicating that SUMOylation efficiency in cells is not dictated by additional effectors, and instead is a direct and intrinsic property of the ligand-bound ERα complex. Collectively, our cellular and *in vitro* studies establish a hierarchical mechanism of SERD activity, in which PIAS ligases directly recognize and SUMOylate SERD-bound ERα. Ubiquitylation by STUbLs is independent of SERD binding yet strictly requires prior SUMOylation. Hence, SERDs do not directly induce ERα ubiquitylation but functionalize a native ERα:PIAS association to enhance SUMOylation, a mechanism that clearly differentiates them from conventional small-molecule degraders such as proteolysis-targeting chimeras (PROTACs) or molecular glue degraders.

We next turned to investigating the structural mechanisms underlying SERD-based ERα SUMOylation. ERα agonists and antagonists typically induce distinct helix12 (H12) conformations, facilitating the recruitment of co-activators or co-repressors, respectively (Fig S6A). A Fulvestrant-bound ERα structure has not yet been resolved, however, the Fulvestrant derivate ICI 164,384 triggers complete H12 dissociation in rat ERβ, providing a possible structural basis for SERD-dependent ERα SUMOylation and degradation (*6, 43, 44*). Other crystal structures of ERα bound to SERDs, including Giredestrant, are based on ERα harbouring stabilizing mutations that aid crystallization (*45, 46*) but prevent SERD-dependent SUMOylation (*7*). We thus attempted to solve the structures of SERD-bound ERα free of mutations that might interfere with SUMOylation. To this end, we devised a BRIL fusion tag (*47*) in combination with a *de novo* designed ERα binder (*48*) compatible with ERα SUMOylation (fig. S6B) and resolved cryo-EM structures of Fulvestrant- and Giredestrant-bound ERα^LBD^ at 3.2 and 3.5 Å, respectively (Fig. 2G-I, fig. S4, fig. S5, fig. S6C-F, Table S3). Our ERα:Fulvestrant structure recapitulates the reported H12 destabilization, with the hydrophobic tail of Fulvestrant protruding into the co-activator binding groove, where it occupies a space incompatible with known agonist- or antagonist states (Fig. 2G-I and fig. S6D), consistent with the complete loss of co-activator recruitment observed in proximity profiling and *in vitro* binding assays (Fig. 2D, fig. S3E-H). In line with previous structures of SUMOylation-deficient ERα mutants, Giredestrant did not induce H12 unfolding, even though it promoted ERα SUMOylation and degradation with similar efficiency as Fulvestrant (fig. S3M and N). Instead, Giredestrant-bound ERα adopted a classical antagonistic H12 conformation virtually indistinguishable from 4-OHT, with H12 occluding the co-activator binding groove (1.5 Å RMSD over 198 residues; fig. S6G). Together, these structures indicate that SERD-based SUMOylation and degradation are not triggered by an antagonistic H12 conformation or unfolding, instead suggesting a critical role for ERα regions beyond the LBD in coupling SERD binding to ERα SUMOylation.

We thus used *in vitro* SUMOylation assays to delineate the involved protein domains. For PIAS4, we found the N-terminal SAP domain dispensable but the PINIT domain and C-terminal SUMO interacting motif (SIM) essential (fig. S6H and I). For ERα, we observed SERD-dependent SUMOylation with intact full-length ERα as well as with a truncated version harboring only the DNA-binding- and ligand-binding domains (DBD-LBD), but not with isolated LBD alone (fig. S6J). Although DNA was not required for SUMOylation (fig. S6K), this suggested a critical role for the ERα DBD. Increasing amounts of isolated LBD progressively reduced SUMOylation of full-length ERα *in trans* (fig. S6L), presumably by competing for access to the SUMOylation machinery despite being insufficient for productive SUMOylation. Together, our structural and biochemical data suggest that SERD-induced ERα antagonism and SUMOylation are structurally uncoupled processes, with the DBD, alongside the LBD, playing a critical role in promoting ERα SUMOylation and subsequent degradation by STUbLs.

### ERα degradation compromises SERD efficacy

Even though induced ERα degradation is widely accepted as the mechanism underlying full antiestrogenic activity of SERDs (*14–17, 49*), some previous reports suggest that degradation may not be causative for their activity (*8, 11*). Having deciphered the ERα degradation pathway triggered by SERDs, we could thus, for the first time, apply genetic loss-of-function studies to directly test the importance of ERα degradation for the therapeutic efficacy of SERDs. In an MCF7 xenograft model, 3xRNF KO cells remained fully sensitive to Fulvestrant treatment despite impaired ERα degradation (fig. S7A). Fulvestrant treatment resulted in loss of cell proliferation (Fig. 3A, fig. S7B) and significant tumor shrinkage (Fig. 3B, fig. S7C and D) independent of ERα degradation, clearly indicating that degradation is not essential for SERD efficacy *in vivo*.

**Fig. 3.**
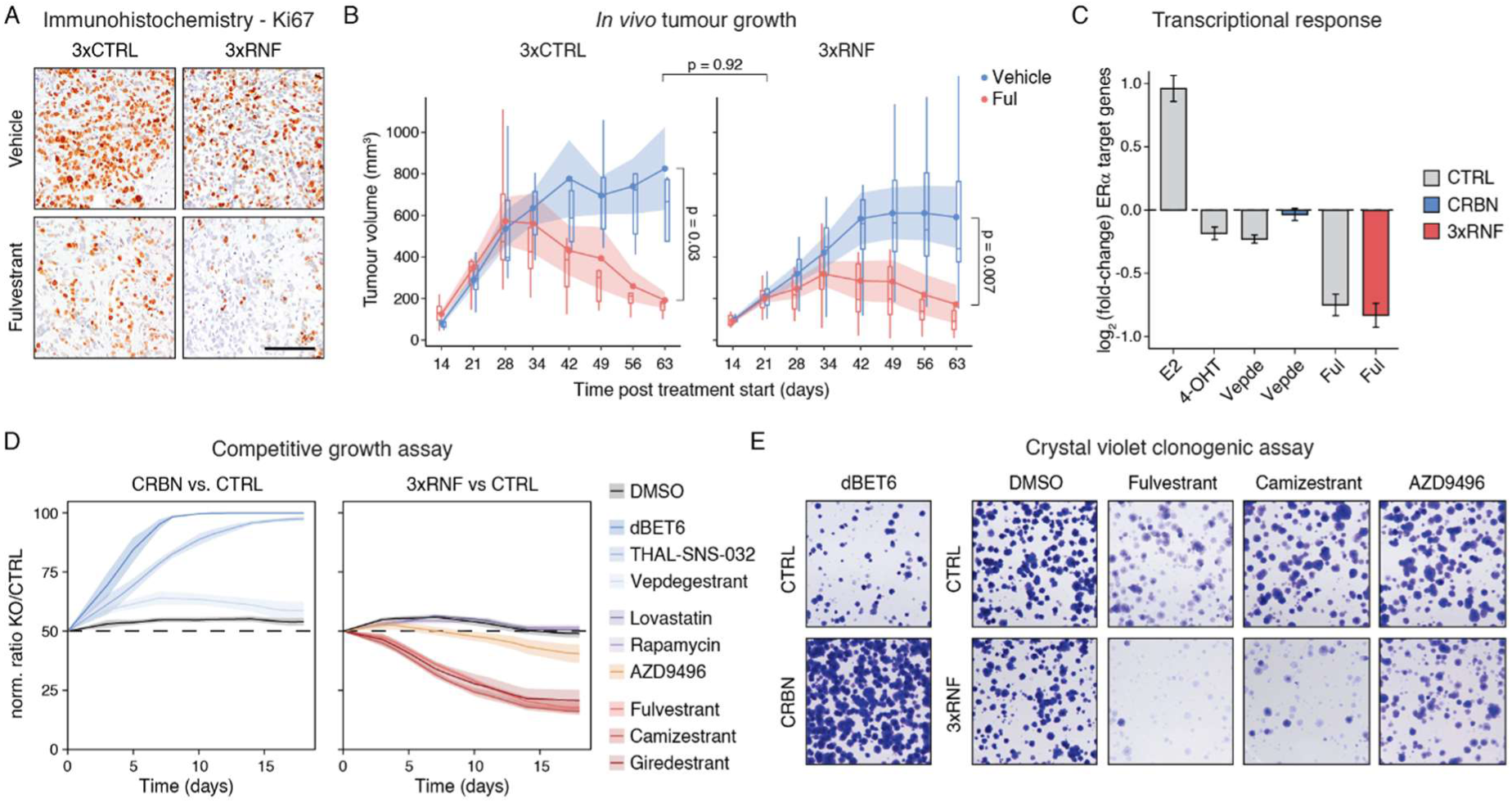
ERα degradation counteracts SERD efficacy. **(A)** *In vivo* tumor cell proliferation. MCF7 iCas9 control or 3xRNF knockout cells were injected into nude mice mammary glands. After tumors were established, mice were subcutaneously treated with two doses of vehicle or Fulvestrant (150 mg/kg) before tumors were immunohistochemically stained for Ki67 expression (brown). Blue, hematoxylin counterstaining; scale bar, 100 μm; images show representative regions from n = 2 (3xCTRL + vehicle) or 3 (all others) tumors per group; Quantification, fig. S7B. **(B)** *In vivo* tumor growth. MCF7 xenografts were established as in (A) and vehicle or Fulvestrant were administered subcutaneously once weekly. Tumor size was measured at indicated timepoints after cell injection. Response by genotype, Welch’s two-sample t-test on log-transformed area under the growth curve. Differential response between genotypes, two-way ANOVA. Points and ribbons, Mean ± SEM of n at start of treatment = 8 (CTRL vehicle), 10 (CTRL Fulvestrant), 7 (3xRNF vehicle), 10 (3xRNF Fulvestrant) mice; boxes, interquartile range (IQR) and median; whiskers, 1.5 x IQR; Growth curves for individual mice, fig. S7C; %Tumor growth inhibition, fig. S7D. **(C)** Transcriptional profiling of ERα activity. MCF7 iCas9 control (gray), CRBN knockout (blue) and 3xRNF (red) knockout cells were treated with DMSO or Estradiol, 4-OHT, Fulvestrant (all 10 nM) or Vepdegestrant (100 nM) for 24 h before transcriptional profiling *via* QuantSeq. Estradiol-treated cells and respective DMSO control cells were starved of hormones for 2 days prior to treatment. n = 3 biological replicates, mean ± SEM of 200 ERα response genes (MsigDB Hallmark ESTROGEN_RESPONSE_EARLY) (*50*). **(D)** Competitive growth assay. GFP expressing MCF7 iCas9 control cells were mixed with mCherry expressing CRBN or 3xRNF knockout cells and treated with DMSO, dBET6 (250 nM), CDK9-degrader THAL-SNS-032 (25 nM), Vepdegestrant (50 nM), AZD9496 (0.5 nM), Rapamycin (2.5 μM), Lovastatin (25 μM), Fulvestrant (0.5 nM), Camizestrant (0.25 nM) or Giredestrant (0.25 nM). Percentage of mCherry^+^ cells was evaluated in regular intervals via flow cytometry. Lines and ribbons, mean ± SEM for n = 7 (DMSO, Fulvestrant, Giredestrant), 5 (AZD9496), 4 (Vepdegestrant) or 3 (all others) independent experiments. **(E)** Colony formation assay. MCF7 iCas9 control, CRBN or 3xRNF knockout cells were treated with DMSO, Fulvestrant (0.5 nM), Camizestrant (0.25 nM), AZD9496 (0.5 nM) or dBET6 (250 nM) for 13 days before staining with crystal violet. Images representative of n = 3 independent experiments with 3 biological replicates each; quantification, fig. S7K.

To decouple the effects of degradation on transcriptional signaling and long-term viability, we next evaluated SERD efficacy in cellular systems. As expected, Estradiol induced the expression of known ERα target genes while 4-OHT and Fulvestrant repressed these genes (Fig. 3C, fig. S7E and F), even though DNA binding affinity remained unaffected by either compound (fig. S7G). The recently FDA-approved ERα PROTAC Vepdegestrant (Arv-471) showed transcriptional ERα inhibition comparable to 4-OHT in WT cells, but - consistent with the expected degradation-based mechanism - lost its efficacy upon KO of the ubiquitin E3 substrate receptor CRBN (Fig. 3C). In contrast, Fulvestrant invariably blocked ERα transcriptional activity in 3xRNF KO and control (3xCTRL) cells. Together, these results show that SERD-based inhibition of ERα-driven transcription occurs independent of ERα degradation. In line with this observation, Vepdegestrant outperformed all SERDs in Erα degradation (fig. S7H) but only moderately inhibited ERα signaling (Fig. 3C) and proliferation in MCF7 cells (fig. S7I).

In flow cytometry-based competitive growth assays, PROTACS targeting BRD4 (dBET6), CDK9 (THAL-SNS-032) or ERα (Vepdegestrant) for CRBN-based degradation lost their efficacy upon CRBN KO, resulting in rapid outgrowth of these KO cells from mixed cultures (Fig. 3D). SUMO-inducing SERDs Fulvestrant, Camizestrant and Giredestrant, in contrast, showed markedly enhanced efficacy in 3xRNF KO compared to 3xCTRL cells, opposite to the resistance phenotype expected for degraders. The activity of a panel of cytostatic anticancer drugs that induce cell cycle arrest independent of ERα signaling was not affected by the 3xRNF KO. Likewise, the efficacy of AZD9496, a SERD that boosts endogenous ERα turnover without induced SUMOylation (*22*), was unchanged by 3xRNF KO. Additionally, 3xRNF KO resulted in sensitization to SUMO-dependent SERDs only in ERα^+^ MCF7 cells but not in ERα^-^RKO colon carcinoma cells (fig. S7J), together highlighting that the observed sensitization is an ERα-dependent on-target effect rather than unspecific hypersensitivity of 3xRNF KO cells. In colony formation assays, MCF7 cells displayed similar resistance to PROTACs upon CRBN KO but enhanced sensitivity to SUMO-inducing SERDs after 3xRNF KO (Fig. 3E and fig. S3K) and in CellTiter-Glo (CTG) cell viability assays, CRBN KO conferred resistance to PROTACs whereas 3xRNF KO cells remained sensitive to SERDs (fig. S3L). Together, these data highlight that induced ERα protein degradation is not required for functional ERα antagonism and cytostatic SERD activity. On the contrary, impaired degradation partially sensitized breast cancer cells to SERD treatment, indicating that STUbL-dependent degradation of SUMOylated ERα protects breast cancer cells from cytostatic SERD effects. This suggests that SUMOylated ERα exerts a dominant-negative function that represses estrogen signaling and breast cancer cell growth more effectively than induction of ERα loss-of-function via inhibition or degradation.

### SERD-induced ERα SUMOylation recruits transcriptional repressors

In transcriptional profiling and cell viability assays in MCF7 cells, SERDs were generally more effective than 4-OHT and other SERMs (Fig. 3C and fig. S7E-F and I). Since this enhanced efficacy was independent of ERα protein degradation, we turned to search for other effectors that are required for the superior performance of SERDs compared to SERMs. To this end, we performed genome-wide CRISPR viability screens in MCF7 cells to identify knockouts that mediate resistance to SERD treatment. Mirroring the results of our FACS-based screens geared to find effectors of ERα degradation, we found knockouts of the core SUMOylation enzymes SAE1, UBA2 and UBE2I enriched in the cells that survived 14 days of Fulvestrant treatment (Fig. 4A). To validate these effects in single-well assays, we conducted competitive co-culture assays comparing GFP-labelled *AAVS1* CTRL KO cells with mCherry-labelled cells harboring target gene KOs. Core SUMOylation genes are essential for cellular fitness in MCF7 cells (*51*) under normal culture conditions and hence gradually disappeared from the mixed culture over time, akin to the essential control genes *UBA1* and *UBE2M* (Fig 4B). In contrast to basal culture conditions, KO of the core SUMOylation enzymes SAE1 and UBE2I conferred a marked survival benefit in the presence of Fulvestrant. In contrast, KO of the essential UPS components UBA1 and UBE2M did not rescue the cytostatic response induced by Fulvestrant. The protective effect mediated by KO of SUMOylation enzymes was furthermore specific to SERDs, as no survival benefit was observed in response to the CDK4/6 inhibitor Palbociclib, an approved cytostatic drug for ER^+^ breast cancer. In sum, these findings identify active SUMOylation as a prerequisite for the full cytostatic activity of Fulvestrant. Together with our biochemical reconstitutions, this points to a mechanism of SERD action, in which SUMOylated ERα exerts a dominant negative effect that actively represses breast cancer cell growth

**Fig. 4.**
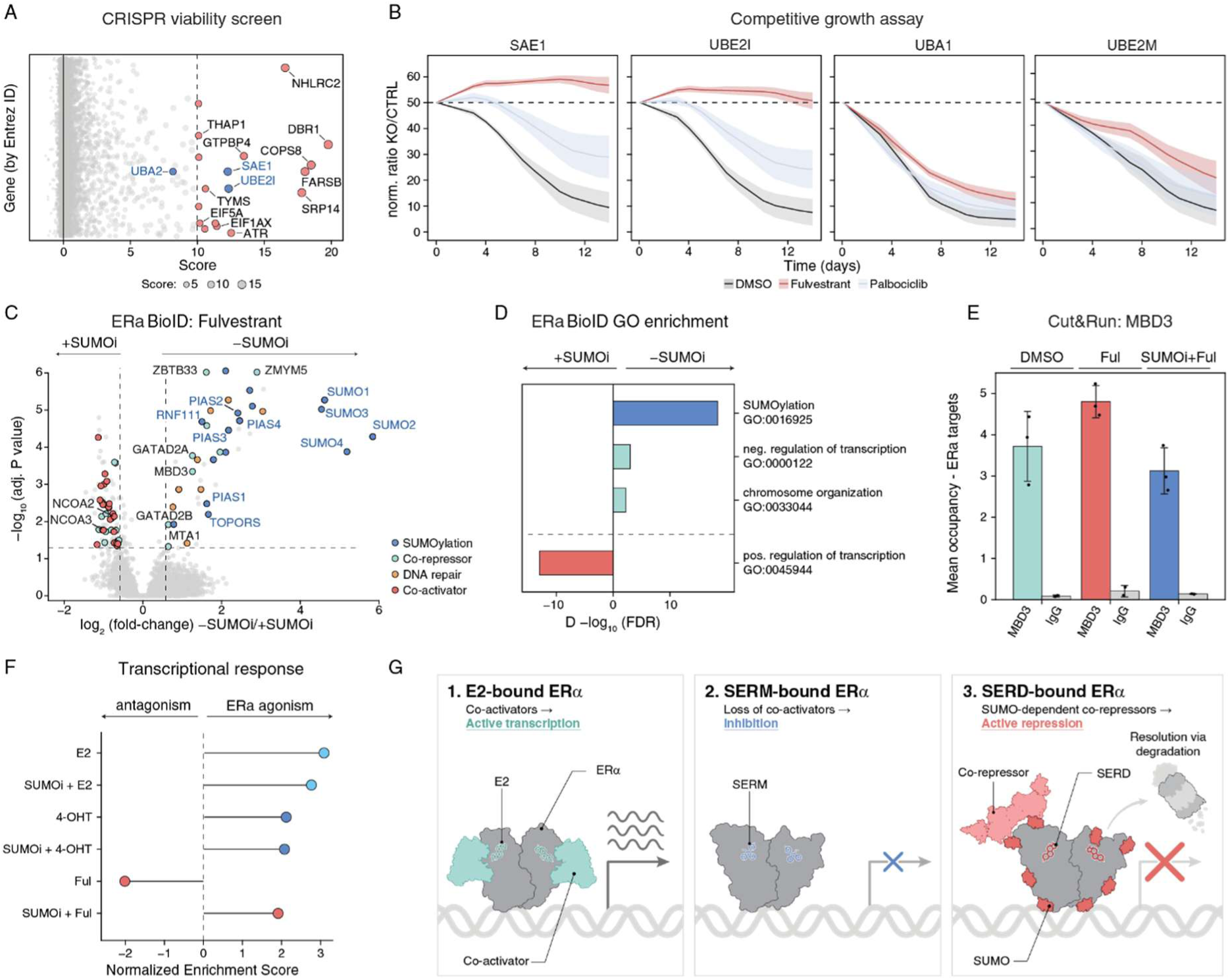
SERD-induced SUMOylation suppresses ERα activity. **(A)** Fulvestrant resistance CRISPR viability screen. MCF7 iCas9 cells were mutagenized with a genome-wide sgRNA library and cultivated in the presence of DMSO or Fulvestrant (10 nM) for 14 days. One-sided MAGeCK analysis of Fulvestrant vs. DMSO, n = 3 biological replicates. Gene-level enrichment score was calculated as -log_10_ (P value) x log_2_ (fold-change). Gene knockouts enriched in Fulvestrant-treated compared to DMSO control population are highlighted in red, core SUMOylation enzymes shown in blue. **(B)** Competitive growth assay. MCF7 iCas9 cells expressing sgRNAs targeting indicated genes alongside mCherry were mixed with cells expressing AAVS1-targeting sgRNA alongside GFP. Cell pools were treated with DMSO, Fulvestrant (5 nM) or Palbociclib (200 nM) and percentage of mCherry^+^ cells was quantified in regular intervals for 14 days via flow cytometry. sgRNA-positive fractions were normalized to day 0. Lines and ribbons, mean ± SEM of n = 6 (DMSO, Fulvestrant) or 5 (Palbociclib) independent experiments. **(C)** ERα BioID proximity labelling. ERα-miniTurbo cells were treated with Fulvestrant (10 nM) with or without ML792 (SUMOi; 2 µM) pre-treatment. Transcriptional activators (red; GOBP:0045944), repressors (cyan; GOBP:0000122), SUMO pathway components (blue; GOBP:0016925) and DNA repair genes (orange; GOBP:0006281, GOBP:0006302) in the scoring window (|FC| > 1.5, adj. P value < 0.05, one-way ANOVA with Tukey’s post hoc test and Benjamini-Hochberg correction, n = 3 biological replicates) are highlighted. **(D)** GO analysis of ERα BioID interactors. Top GO biological process terms overrepresented among ERα interactors upon Fulvestrant treatment in the presence versus absence of SUMOylation inhibition (Δ - log_10_ (FDR); Benjamini-Hochberg-corrected two-tailed Fisher’s exact test). **(E)** CUT&RUN of MBD3 at ERα binding sites. Mean spike-in–normalized MBD3 and IgG signal at Fulvestrant-specific ERα peaks per treatment condition. Bars and error bars, mean ± SD across biological replicates; individual replicate values overlaid as points. **(F)** Transcriptional profiling of ERα activity. MCF7 wild-type cells were hormone-deprived for 3 days, followed by 24 h treatment with Estradiol, 4-OHT or Fulvestrant (all 10 nM) with or without ML792 (SUMOi; 2 µM) pre-treatment for 1 h, and ERα activity was assessed by RNA-Seq of n = 3 biological replicates. Gene set enrichment analysis was performed based on the Hallmark Estrogen Response Early gene set (MSigDB) (*50*), normalized enrichment scores are shown. **(G)** Schematic representation of SERD mechanism. In addition to co-activator displacement caused by SERMs (middle panel), SERD-based ERα SUMOylation recruits co-repressors that actively silence ERα target genes and thereby disrupt estrogen signaling more thoroughly than complete ERα loss-of-function. STUbL-based degradation of SUMOylated ERα resolves active repression and thereby lowers rather than drives ERα efficacy. Ful, Fulvestrant; E2, Estradiol; 4-OHT, 4-Hydroxytamoxifen; LBD, ligand binding domain; H12, Helix12; SUMOi, ML792; GO, gene ontology.

To resolve the underlying molecular cause, we used BioID to profile changes in the Fulvestrant-mediated ERα interactome in the presence versus absence of SUMOylation. As expected, Fulvestrant-treated ERα was associated with diverse SUMO variants, E3 SUMO ligases and STUbLs and this recruitment of the SUMOylation machinery was lost upon SUMOi treatment (Fig. 4C). In SUMO-competent cells, we found ERα interactors enriched in DNA repair proteins and repressive transcriptional co-factors, most notably the nucleosome remodeling and deacetylation (NuRD) complex (Fig. 4C and D). In comparison, SUMOi treatment prevented the recruitment of co-repressors and instead shifted the interactome towards transcriptional activators, with co-activators such as NCOA2 and NCOA3 or transcription factors such as TFAP4 and JUNB associated with ERα specifically in the absence of SUMOylation. Using NanoBiT complementation, we confirmed that Fulvestrant treatment induced a SUMO-dependent interaction between ERα and the NuRD complex subunits HDAC1 and MTA2 (fig. S8A). Accordingly, we observed an enrichment of the NuRD complex subunit MBD3 on ERα occupied sites across the genome after Fulvestrant treatment via CUT&RUN sequencing and, again, this recruitment was absent after the inhibition of SUMOylation (Fig. 4E and fig. S8B and C). Hence, SERD-induced SUMOylation directly drives the recruitment of transcriptional repressors, converting ERα bound chromatin loci from a transcriptionally competent to a fully repressed state.

4-OHT and other SERMs are known to only partially antagonize ERα, and after hormone deprivation, in endometrium or bone, they can even induce ERα activity (*52–54*). In comparison, Fulvestrant and other SERDs are described as pure antagonists devoid of partial agonist activity (*55*). To examine to what extent the SUMO-dependent, repressive ERα interactome contributes to the superior transcriptional disruption of SERDs compared to SERMs, we turned to transcriptional profiling of MCF7 cells via RNA-seq. After hormone deprivation, Estradiol strongly activated ERα activity as expected. We also observed upregulation of ERα target genes by 4-OHT treatment, in line with its reported partially agonistic activity (*52–54*). In comparison, Fulvestrant further repressed ERα even under hormone depletion (Fig. 4F, fig. S8D-F). After SUMOi pre-treatment, the full transcriptional antagonism elicited by Fulvestrant was converted to mild ERα agonistic transcriptional activity comparable to 4-OHT, validating that induced ERα SUMOylation is required for the strong transcriptional antagonism mediated by SERDs.

Together, our data identifies induced SUMOylation as the direct biochemical consequence of SERD binding and highlights SUMOylation as the defining feature of SERD activity that drives their unique fully anti-estrogenic potential. Proteasomal degradation, in contrast, is a secondary consequence of induced SUMOylation that does not drive but rather counteract SERD efficacy by eliminating the dominant negative repression activity of SUMOylated ERα (Fig. 4G).

## Discussion

Selective estrogen receptor degraders derive their name from their capacity to induce proteolytic degradation of ERα. For over two decades, these agents have served as a cornerstone of treatment for advanced breast cancer, widely prescribed due to their superior therapeutic efficacy compared to SERMs. Despite their widespread clinical use, the precise molecular mechanism of SERD-induced ERα degradation and its contribution to the efficacy of this class of medicines has remained elusive.

In this study, we coupled orthogonal ERα stability and SERD resistance CRISPR/Cas9 screens with proximity labelling and biochemical reconstitutions to determine genetic networks required for SERD-induced ERα degradation and cytostatic effects. Using fully reconstituted *in vitro* SUMOylation assays, we establish that PIAS SUMO E3 ligases directly recognize and modify the SERD-bound receptor. This biochemically decouples ligand-induced SUMOylation from the cellular environment and demonstrates that SERDs function by directly remodeling the native association of ERα with its SUMO E3 ligases. The MoA of SERDs is hence functionally differentiated from proximity-inducing drugs such as PROTACs or molecular glues. SERD-induced SUMOylation of ERα promotes recruitment of transcriptional repressors including the NuRD complex, thereby converting ERα from an activating transcription factor into the focal point of a repressive transcriptional assembly. SUMO-dependent repression silences ERα activity more effectively than hormone withdrawal, presumably by inhibiting both ligand-dependent and -independent ERα transcriptional activity (*56, 57*), thus providing a mechanistic explanation for the full antagonism elicited by SERDs compared to SERMs.

In contrast, ERα degradation by the STUbLs RNF4 and RNF111 is a secondary consequence of SERD-induced SUMOylation that does not contribute to ERα repression. Counterintuitively, stabilizing the SERD-bound, SUMOylated ERα through genetic E3 ligase inactivation further enhanced rather than diminished SERD efficacy, likely by increasing the persistence of the repressive environment. This clearly shows that cellular turnover of SUMOylated ERα actively limits, rather than drives, the full potency of SERDs. Our observations are incompatible with a degradation-centric model of SERD activity and instead reveal a mechanism of action wherein SUMOylated ERα recruits co-repressors to actively silence ERα target genes. SUMOylation thus conveys a ‘dominant-negative’ neo-function to ERα, mediating an even stronger functional disruption than complete target elimination.

Target protein SUMOylation, STUbL-mediated ubiquitylation and proteasomal degradation have been reported for clinical compounds beyond SERDs, including approved drugs targeting PARP1/2, DNMTs and TOP1/2, as well as investigational inhibitors of WRN and FEN1 (*58–68*). Resistance to PARP inhibitors partially occurs through genetic PARP1 inactivation (*69, 70*), hinting at a similar dominant-negative pharmacology beyond inhibition as we discovered for SUMO-inducing SERDs. In analogy to SERDs, the PARP1/2 but also DNMT, TOP1/2 and FEN1 drugs are first and foremost inhibitors yet appear to exert additional effects through target protein SUMOylation.

Previous studies have described SERD-induced nuclear ERα immobilization as a proxy for cellular efficacy (*11, 71, 72*). While PARP inhibitors mechanistically trap PARP on chromatin (*66, 73*), ERα immobilization in the nucleus does not appear to be a direct biochemical consequence of SERD-induced ERα DNA binding (fig. S7G). Dissecting the molecular interplay between SUMOylation, target immobilization and gene repression remain priorities for future research.

SUMOylation motifs have been identified as the predominant repressive signal in transcription factors, occurring in ∼29% of known repressive domains (*74*). Combined with the mechanistic dissection of SERD action presented herein, this highlights opportunities and clinical tractability for a new class of drugs that induce SUMOylation-dependent functional target inhibition. While the identification of Fulvestrant and other clinically relevant SERDs as SUMO-inducing drugs was retrospective, increasing our mechanistic understanding of ligand-triggered target SUMOylation will empower the purposeful molecular design of a therapeutic modality geared to target difficult-to-drug transcription factor

## Acknowledgements

The authors thank all members of the Thomä and Winter laboratories for experimental advice and helpful feedback; L. Haas and M. Schütz-Stoffregen for help with manuscript editing; J. Carvalho for support on graphics design; the Core Facility Flow Cytometry of the Medical University of Vienna and the BioOptics facility of the IMP/IMBA/GMI for flow cytometry support; the CeMM Biomedical Sequencing Facility for NGS sample processing and sequencing; J. Zuber at the Research Institute of Molecular Pathology for sharing iCas9 cell lines, plasmids and sgRNA libraries; A.-C. Gingras for providing BioID plasmids; R. Hay for sharing an anti-RNF4 antibody; A. P. Kutschat for Cut&Run advice; F. Ö. Elaslan for advice on *ex vivo* tissue dissociation; N. Mailand and P. Haahr for helpful discussions.

## Author contributions

M.H., C.S. and S.S. contributed equally and will be putting their name first on this work on their respective CVs. M.H., C.S., S.S., N.H.T. and G.E.W. conceived and planned this project. M.H., C.S., S.S. and Ma.S. designed and conducted experiments with help from I.K., My.S. and K.K.; M.H., C.S., S.S. and Ma.S. performed data analysis and generated figures. Ma.S. and I.K. performed *in vivo* experiments. S.S., L.K., G.K. and S.C performed structural analysis. F.F. performed BioID data analysis. H.I. performed CRISPR screen analysis. D.H., P.B., M.M., M.P and J.D.A. designed critical reagents and established experimental workflows. S.H. analyzed IHC data. D.S., B.E.C and A.C.O. supervised research and provided scientific input. M.H., C.S., S.S., N.H.T. and G.E.W. co-wrote the manuscript with input from all co-authors.

## Data, code and materials availability

The mass spectrometry proteomics data have been deposited in the ProteomeXchange Consortium (*75*) via the PRIDE partner repository (*76*) with the project DOI 10.6019/PXD069276. RNA-seq, 3′ mRNA-seq (QuantSeq) and CUT&RUN raw and processed data have been deposited in the NCBI Gene Expression Omnibus under accession codes GSE323330, GSE323374 and GSE336383, respectively. The cryo-EM density maps of tetrameric BRIL-ERα^LBD^ in complex with the ERα^LBD^ binder bound to Fulvestrant have been deposited in the Electron Microscopy Data Bank under accession codes EMD-58680 (local refinement map of dimer^1^), EMD-58681 (local refinement of dimer^2^), EMD-58683 (local refinement of binder), EMD-58679 (consensus map), EMD-58678 (composite map) and for Giredestrant under accession code EMD-58682. The corresponding atomic coordinates for the Fulvestrant-bound- and Giredestrant-bound complex have been deposited in the Protein Data Bank under accession code 31VT and 31VU, respectively. All datasets will be made publicly accessible upon acceptance of the peer-reviewed publication. Raw imaging data from colony formation assays and IHC staining are available from the corresponding authors upon reasonable request.

All biological materials generated in this study are available from the corresponding authors upon reasonable request and under material transfer agreements with the respective host institutions (AITHYRA Research Institute for Biomedical Artificial Intelligence of the Austrian Academy of Science, CeMM Research Center for Molecular Medicine of the Austrian Academy of Sciences, or Friedrich Miescher Institute for Biomedical Research), as appropriate.

Custom scripts used for image analysis (ImageJ macros and Python scripts) are available from the corresponding authors upon reasonable request. All other analyses were performed using previously published pipelines and publicly available software as described in the Methods.

## Supplementary materials

### Materials and Methods

#### Plasmids and oligonucleotides

The design and construction of the human genome-wide sgRNA library (*77*) used for ERα stability screens in KBM7 cells and viability-based CRISPR screens in MCF7 cells, the UPS-focused sgRNA library (*78*) used for ERα stability screens in MCF7 cells, lentiviral sgRNA expression vectors used for single gene knockouts, and lentiviral vectors used for the engineering of inducible Cas9 cell lines have been described previously (*77, 79*). Lentiviral vectors used for dual gene knockouts were modified from Dual-sgRNA_hU6-mU6 (*80*) (Addgene plasmid #154194; gift from Anna Obenauf) and vectors for simultaneous triple gene knockout were obtained by introducing a bovine U6-filler-tracrRNA array synthesized as gene fragment (Twist Bioscience) upstream of the dual sgRNA expression cassette while the restriction sites for all three guides were changed to BsmBI to allow simultaneous cloning of all sgRNAs via BsmBI-v2 Golden Gate Assembly Kit (NEB #E1601). The ERα-GFP-P2A-mCherry and ERα-GFP-P2A- miRFP670nano3-H2A protein stability- and immobilization-reporters, respectively, were obtained by modifying a BRD4 stability reporter described previously (*81*).

The dox-inducible pSTV6_ERα_3xFlag_miniTurbo and pSTV6_NLS-GFP_3xFlag_miniTurbo plasmids were generated using the gateway cloning (BP Clonase II, 11789020 and LR Clonase II, 11791100 both Invitrogen) donor vectors pENTR221_ERα and pENTR223_NLS-GFP and the destination vector pSTV6_ccdB_ 3xFlag_miniTurbo (provided by A. Gingras).

For the generation of the ERα-GFP-P2A-mCherry reporter, ERα was cloned into a pRRL lentiviral vector, fused to mEGFP and linked to mCherry via a self-cleaving P2A peptide for normalization, using HiFi assembly (NEB #E2621L).

Split-HaloTag plasmids were derived from pCDNA5/FRT-Lyn11-mEGFP-cpHaloΔ-(GGS)9-FKBP-P2A-Hpep3-(GGS)3-FRB-mScarlett vector (Addgene plasmid #205703; a gift from Kai Johnsson). EGFP-cpHaloΔ and Hpep3-(GGS)3 were individually cloned into pRRL lentiviral backbones to derive pRRL_SFFV-EGFP-cpHaloΔ-(GGS)9-ERα and pRRL_SFFV-Hpep3-GGGS-SUMO3-EF1α-TagBFP used to quantify ERα SUMOylation.

The ERα-GFP-P2A-H2A-IRFP reporter plasmid used for chromatin FACS experiments was generated by modification of the parental ERα-GFP-P2A-mCherry reporter. The mCherry sequence was replaced by H2A (taken from Addgene plasmid #158691, a gift from Ravid Straussman) and miRFP670nano3 using HiFi assembly.

If possible, number of sgRNAs were matched between control and test knockouts, i.e. single KOs were compared to a single neutral control sgRNA (AAVS1), combinatorial RNF4 and RNF111 KO was compared to a combination of two neutral sgRNAs (AAVS1_3 and AAVS1_4) and combinatorial RNF4, RNF111 and RNF216 (3xRNF) KO was compared to combinatorial knockout of three neutral control genes (AAVS1, OR51T1, LALBA; 3xCTRL).

All plasmids used in this study are shown in Table S1, all oligonucleotides, sgRNAs and shRNAs in Table S2.

#### Molecular cloning of sgRNA combinations

For cloning of dual or triple sgRNA combinations, individual sgRNAs were synthesized as single stranded DNA oligos (Microsynth) with unique overhangs specifying the respective position in the dual or triple sgRNA vector. Single stranded oligos were annealed and phosphorylated as described previously, and cloned into dual or triple sgRNA recipient plasmids simultaneously in a single cyclic digest-ligation reaction using BsmBI-v2 NEBridge® Golden Gate Assembly Kit (NEB #E1602) according to manufacturer’s instructions.

#### Cell culture

Human MCF7 cells, originally sourced from ATCC (HTB-22), and Lenti-X 293T lentiviral packaging cells (Takara, #632180) were cultured in DMEM (Sigma-Aldrich) supplemented with 10% fetal bovine serum (FBS; Thermo Fisher), 100 U/mL penicillin/streptomycin (Sigma-Aldrich), 2 mM L-glutamine (Gibco), 1 mM sodium pyruvate (Gibco) and MEM Non-Essential Amino Acids Solution (1x; Gibco). MCF7 iCas9 cells were generated as previously described (*79*). Human KBM7 iCas9 cells, gifted from Johannes Zuber (Research Institute of Molecular Pathology), were cultured in IMDM (Sigma-Aldrich) supplemented with 10% FBS, penicillin/streptomycin, L-glutamine and sodium pyruvate as above. Human RKO iCas9 cells, gifted from Johannes Zuber (Research Institute of Molecular Pathology), were cultured in RPMI 1640 (Sigma-Aldrich) supplemented with 10% FBS, penicillin/streptomycin, L-glutamine and sodium pyruvate as above. Human T47D cells (Cytion, #300353) were cultured in RPMI 1640 supplemented with 10% FBS, penicillin/streptomycin, L-glutamine, sodium pyruvate and MEM Non-Essential Amino Acids as above. Human EFM-19 (DSMZ, #ACC 231) and CAMA-1 cells (ATCC, #HTB-21) were cultured in DMEM/F12 medium supplemented with 10% FBS, penicillin/streptomycin, L-glutamine, sodium pyruvate, MEM Non-Essential Amino Acids and 10 μg/mL insulin (Sigma-Aldrich, #I9278). All cell lines were authenticated by STR profiling and routinely tested for mycoplasma contamination.

#### Lentivirus production and transduction

Semiconfluent Lenti-X cells were co-transfected with lentiviral plasmids, the lentiviral pCMVR8.74 helper (Addgene plasmid #22036) and pMD2.G envelope (Addgene plasmid #12259; both gifts from Didier Trono) plasmids using polyethylenimine (PEI) transfection (PEI MAX® MW 40,000, Polysciences). Virus containing supernatant was clarified by centrifugation. Target cells were infected at limiting dilutions in the presence of 4 μg/mL polybrene (Santa Cruz Biotechnology).

### BioID proximity labelling and quantitative mass spectrometry

#### Cell culture and treatment

BioID experiments were performed in doxycycline-inducible NLS-GFP-miniTurbo or ERα-miniTurbo expressing MCF7 cells. Cells were seeded in 10 cm dishes and induced with doxycycline (1 µg/mL) for 24 h prior to treatment. All samples were pre-treated with 1 µM Carfilzomib for 1 h, while half of the samples received additionally 2 µM ML792. Samples were treated with either 10 nM 17β-Estradiol (E2) or 10 nM Fulvestrant for 6 h. All samples were supplemented with 50 µM biotin for the final hour of treatment. Following treatment, cells were washed with PBS, trypsinized, pelleted, washed again with PBS, and snap-frozen.

#### Lysis, enrichment and on-bead digestion

Cell pellets were lysed with 200 µL lysis buffer (1% SDS, 2 mM MgCl2 in PBS) supplemented with 1x Halt protease inhibitor (Thermo Scientific, 11804111) and benzonase (Merck, 70746). Lysates were incubated at 37 °C for 30 min with intermittent vortexing, clarified by centrifugation (18,000 g, 30 min, 4 °C) and quantified using the Pierce 660 nm protein assay (Thermo Scientific, 22660) with the Ionic Detergent Compatibility Reagent (Thermo Scientific, 22663). Equal protein amounts (3 mg per sample) were adjusted to 300 µL in lysis buffer and reduced with 4.5 mM TCEP for 1 h at 56 °C. pH was adjusted by adding 200 mM HEPES Ph 7.5. Samples were then alkylated with 20 mM iodoacetamide for 30 min at room temperature (RT). Streptavidin agarose beads (50 µL per sample; Thermo Scientific, 20353) were pre-washed with PBS and added to the lysates for 1 h at RT. Beads were subsequently washed using a vacuum manifold: twice with Wash Buffer 1 (0.2% SDS in PBS), 16 times with Wash Buffer 2 (8 M urea in PBS), and four times with PBS. Beads were resuspended in digestion buffer (50 mM ammonium bicarbonate, 0.2 M guanidine hydrochloride, 1 mM CaCl2 in HPLC-grade H2O) and transferred to protein low-binding tubes. On-bead digestion was initiated by adding 1 µg of sequencing-grade trypsin (Promega, V5117) and incubated overnight at 37 °C.

#### Peptide cleanup, TMT labelling, and fractionation

Peptides were separated from the beads by centrifugation and washed with HPLC-grade H2O. Combined digests were acidified with trifluoroacetic acid (1% final) and subjected to solid-phase extraction using in-house packed C18 stage tips. Peptides were eluted in 90% acetonitrile/0.1% TFA, dried by vacuum centrifugation, and stored at −20 °C. Dried peptides were reconstituted in 100 mM HEPES (pH 8.5) and labelled using TMTpro™ 18plex reagents (Thermo Scientific, A52045) according to the manufacturer’s protocol. Reactions were incubated at RT for 1 h, quenched with 0.385% hydroxylamine, and pooled into a single tube. Pooled peptides were fractionated using in-house packed C18 columns with reversed-phase buffers (20 mM ammonium formate mixed with increasing concentrations of acetonitrile: 16%, 20%, 24%, 28%, 80%). Five fractions were collected, dried, and stored at −20 °C until LC-MS/MS analysis.

#### MS data acquisition

Mass spectrometry analysis was performed on an Orbitrap Fusion Lumos Tribrid mass spectrometer coupled to a Dionex Ultimate 3000 RSLCnano system via a Nanospray Flex Ion Source interface (all components Thermo Fisher Scientific, San Jose, CA). Peptides were loaded on a trap column (PepMap 100 C18, 5 μm, 5 × 0.3 mm, Thermo Fisher Scientific, San Jose, CA) with 0.1% TFA in HPLC-grade H_2_O (1.15333.2500, MERCK) at a flow rate of 10 µL/min. The trap column was switched in-line, and peptides were separated on an analytical column (50 cm, 75 µm inner diameter, packed with ReproSil-Pur 120 C18-AQ, 3 µm, Dr. Maisch, Ammerbuch-Entringen, Germany), fitted with a fused silica (20 μm ID x 7 cm L x 365 μm OD; Orifice ID: 10 μm, CoAnn Technologies) ESI emitter. During the analytic separation the column was kept at 50°C. Peptides were separated over an analytical gradient of 190 min with a constant flow rate of 230 nL/min. As buffer A, 0.4% formic acid (FA, 1.11670.1000, MERCK) in HPLC-grade H2O and for buffer B, 0.4% FA in acetonitrile (ACN, VWR Chemical, 83640.32) was used. The LC gradient was set up as follows: isocratic at 6% buffer B (0-4 min), linear increase to 9% (4-5 min), followed by a linear gradient to 30% by 146 min and a second steeper gradient to 65% buffer B by 154 min. The column was subsequently flushed with 100% organic buffer (buffer B) and re-equilibrated at 6% from 167-190 min. The mass spectrometer was operated in a data-dependent acquisition (DDA) mode using a maximum of 10 dependent scans (TopN approach) with synchronous precursor selection (SPS) enabled. Peptide ionization at a constant voltage of 1.8 kV. MS1 precursor survey scans for MS2 and MS3 levels were acquired with scan range of 400 - 1600 m/z and a resolution of 120,000 (at 200 m/z) in the Orbitrap. For peptide precursor scans, the automatic gain control (AGC) was set to ‘standard’ with a maximum injection time of 50 ms. For precursor ions only charge states of 2-5 were selected, whereas undetermined charge states were excluded. The dynamic exclusion was set to 60 s with a ±10 ppm window and the intensity threshold was set to 5.0e3. For MSn scans, a charge-state filter was used. MS2 spectra were obtained in a dual-pressure linear ion-trap, using a single charge state per branch (z = 2 to z = 5). Ions were isolated using the quadrupole with a ±0.7 m/z isolation window. Fragmentation was achieved by collision-induced dissociation (CID) with a fixed normalized collision energy of 35% and an CID activation time of 10 ms. For MS2 scans, the normalized AGC target was set to 200% with a maximum injection time of 35 ms. For MS3 scans, precursor ions were isolated using SPS waveform with varying isolation windows for charge stats: 1.3 m/z for z = 2, 1.2 m/z for z = 3, 0.8 m/z for z = 4 and 0.7 m/z for z = 5. Fragment ions were further fragmented by high-energy collision induced dissociation (HCD) at a fixed activation energy at 45% collision energy. The AGC target was set to 300% with a maximum injection time of 100 ms. Orbitrap scan range was set to 100 - 500 m/z at a resolution of 50,000.

#### MS data processing

Following data acquisition, MS-raw files were processed using the Proteome Discoverer pipeline tool (PD, Thermo Fisher Scientific, version 2.4.1.15) using a TMTpro 18-plex quantification method. Peptide identification search was performed with Sequest HT, querying for fully tryptic peptides. As parameters, a maximum of two missed cleavages and a peptide length of 6-144 amino acid, with a peptide precursor mass tolerance of 10 and a fragment ion mass tolerance of 0.6 DA respectively. Spectra were searched against the canonical human protein database obtained from UniProtKB (download 05.11.2021, 20,304 sequences). The reference proteome was supplemented with an internally curated list of 298 common lab contaminants, including keratins, bovine serum albumin (BSA), and streptavidin. As variable modification Methionine oxidation (+15.994 Da), Deamidation (0.984 Da), phosphorylation on Serine, Threonine and Tyrosine (+79.966 Da) and N-terminal specific acetylation (+42.011 Da), methionine loss (-131.040 Da) and acetylation with methionine loss (-89.030 Da) with a maximum number of three variable modification of the same type and a total of four modifications per peptide. Carbamidomethylation (+57.021 Da) of cysteine residues and tandem mass tag (TMT) 18-plex labelling of peptide N termini and lysine residues (+304.207 Da) were used as static modification. PSM and peptide FDR were controlled by Percolator at 1% respectively. Obtained results were filtered to include only spectrum matches with a Sequest HT cross-correlation factor (Xcorr) larger or equal to 0.9. Phopshosites needed a minimum site-probability of 75 corresponding to the high confidence threshold. For protein abundance inference, only high confidence proteotypic peptides were included. Protein and peptide quantification was based on TMTpro reporter ion intensities. Where applicable, reporter abundances were derived from signal-to-noise (S/N) ratios; otherwise, raw reporter ion intensities were used. Isotopic impurity correction was enabled. A co-isolation threshold of 80% was applied to remove precursors with too high co-isolation interference. To reduce noise, peptides with an average TMTpro reporter ion S/N ≤ 10 were excluded. Additionally, an SPS mass match threshold of 65% was used to remove peptides with high interference. For further processing the raw protein abundances were exported from proteome discoverer.

#### BioID data analysis

The raw protein abundances of the 3408 proteins obtained from Proteome Discoverer were normalized by total signal normalization and scaled to the median of the total signal across samples. After normalization, proteins flagged as common lab contaminants (33 proteins) were filtered from the dataset. Next, all proteins which showed a coefficient of variation (CV) > 100 within one grouped experimental condition and proteins with low abundant signals (70 proteins) were filtered across the entire experiment. This filtering strategy resulted in a total of 3305 protein identifiers retained in the dataset for further analysis. Next, the abundances per condition were grouped to derive the mean abundances per sample condition. Following this step, the abundances were log_2_-transformed, and the log_2_FC against the GFP controls was calculated for Estradiol (6 h), Ful (6 h) and ML792 + Ful (6 h) treatment. Further, the comparison of Estradiol (6 h) against Ful (6 h) and Ful (6 h) vs. ML792 + Ful (6 h) treatment was included in the analysis. Abundance changes were tested for their significance, by employing a one-way ANOVA (two-sided) on individual proteins across biological triplicates independently for each condition. Post hoc comparisons were conducted using a Tukey’s honest significant difference (HSD) test (two-sided). P values obtained from the Tukey’s HSD tests were corrected for multiple testing with the Benjamini-Hochberg (BH) procedure. The same 18plex analysis also included two additional conditions: Ful (1 h) and ML792 + Estradiol (6 h). These two conditions were subsequently filtered out, as their profiles aligned closely with other tested conditions and did not contribute additional insights.

Significantly differentially expressed genes were defined based on adjusted P value and log_2_FC thresholds: adj. P value < 0.001 and fold-change > 3 in Fig. 2C and D, and adj. P value < 0.05 and |fold-change| > 1.5 in fig. S3E and Fig. 4C. Gene lists were subjected to statistical overrepresentation analysis using PANTHER (v19.0) with GO-Slim annotations (*82*). Fisher’s exact test with false discovery rate (FDR) correction was applied. Overarching biological process (BP) terms were manually curated based on the PANTHER hierarchy, retaining those significant in at least one comparison. Terms showing -log_10_(ΔFDR) values > |2.225| between conditions were visualized. All data analyses were performed in R (version 4.4.1) using RStudio (2025.05.1).

### Cleavage Under Targets and Release Using Nuclease (CUT&RUN)

CUT&RUN was performed as described previously (*83*), with some modifications following the EpiCypher CUT&RUN protocol v2.0. MCF7 wild-type cells were seeded in 10-cm dishes at a density of 4 × 10^6^ cells per dish. The next day, cells were pre-treated with Carfilzomib (1 µM) for 1 h in absence or presence of the SUMOi ML792 (2 µM), followed by treatment with Estradiol or Fulvestrant (each at 10 nM) for 1h.

1 × 10^6^ cells per reaction were harvested and nuclei were isolated by incubation in nuclear extraction buffer (20 mM HEPES pH 7.9, 10 mM KCl, 0.1% Triton X-100, 20% glycerol, 1 mM MnCl_2_, 0.5 mM spermidine, protease inhibitors), followed by binding to activated concanavalin A-coated magnetic beads (EpiCypher, 21-1411). Bead-bound nuclei were incubated with primary antibodies in antibody buffer (digitonin buffer with 2 mM EDTA) overnight at 4°C with gentle agitation. CUT&RUN was carried out using antibodies against ERα (Cell Signaling Technology, #13258), MBD3 (Cell Signaling Technology, #99169) or control IgG (Cell Signaling Technology, #3900) in 3 biological replicates per condition for ERα and MBD3, and 2 biological replicates per condition for IgG. After washing with digitonin buffer (20 mM HEPES, 150 mM NaCl, 0.5 mM spermidine, 0.01% digitonin, protease inhibitors), samples were incubated with 700 pg/mL protein A/G–micrococcal nuclease (pAG–MNase), followed by controlled MNase activation via addition of 2 mM CaCl_2_. Targeted chromatin digestion was performed at 4°C for 2 h, and reactions were stopped by addition of stop buffer (340 mM NaCl, 20 mM EDTA, 4 mM EGTA, 50 µg/mL RNAse A, 50 µg/mL glycogen) supplemented with 0.5 ng *Escherichia coli* (*E. coli*) spike-in (EpiCypher, 18-1401) and incubation at 37°C for 10 min. Released chromatin fragments were recovered from the supernatant after bead separation, and DNA was purified using the Monarch Spin PCR & DNA Cleanup kit (New England Biolabs, T1130L).

DNA yields were quantified and 5 ng DNA was used for sequencing library preparation with the NEBNext Ultra II DNA Library Prep Kit for Illumina (New England Biolabs, E7645L) according to the manufacturer’s instructions. Libraries were amplified by PCR using NEBNext Multiplex Oligos for Illumina (Dual Index Set 1) primers (New England Biolabs, E7600S), purified with Mag-Bind TotalPure NGS beads (Omega Bio-tek, M1378-01), quantified, pooled, and sequenced on an Illumina NovaSeq X Plus platform (paired-end 150 bp).

Sequencing data were processed using the nf-core/cutandrun pipeline (v3.2.2; doi: 10.5281/zenodo.5653535). Reads were aligned to the human reference genome (GRCh38, NCBI assembly) using Bowtie2, with *E. coli* used as a spike-in reference. Coverage tracks were normalized to the *E. coli* spike-in, scaling each sample by a normalization factor derived from its number of *E. coli*-aligned reads to generate spike-in–normalized BigWig files. Peaks were called using MACS2 (v2.2.7.1) at a stringency of q < 0.01, using the matched IgG sample as control, and ENCODE blacklist regions (hg38 v2) were excluded from downstream analyses. A consensus set of Fulvestrant-responsive ERα binding sites was defined as ERα peaks reproducible in at least two of the three biological replicates (Fulvestrant condition). MBD3 CUT&RUN signal was quantified at this consensus peak set using deepTools (v3.5.1) multiBigwigSummary (BED-file mode), extracting mean signal from the spike-in–normalized BigWig files over peak regions. Signals were averaged per replicate and condition, and summary statistics (mean ± SD) were calculated across biological replicates (n = 3 for MBD3; n = 2 for IgG). CUT&RUN signal matrices were computed with deepTools (v3.5.1) computeMatrix in reference-point mode, centered ±5 kb around the consensus Fulvestrant-responsive ERα binding sites with a bin size of 50 bp and skipping regions with zero coverage (--skipZeros), and visualized using plotProfile.

### 3’ mRNA-sequencing (QuantSeq)

MCF7 AAVS1 KO, RNF4/111/216 KO, and CRBN KO cells were seeded in 6-well plates (5 × 10^5^ cells/well) 48 h before treatment. For SERM, SERD and PROTAC comparisons, cells were cultured in DMEM supplemented with 10% FBS and treated for 24 h with DMSO, 10 nM 4-OHT, 10 nM Fulvestrant, or 100 nM Vepdegestrant. For the Estradiol treatment, cells were hormone-deprived for 48 h in phenol red-free DMEM (Thermo Fisher Scientific, 21063029) supplemented with 10% charcoal-stripped FBS (Thermo Fisher Scientific, A3382101) and subsequently treated with 10 nM Estradiol for 24 h. For the harvest cells were washed with PBS, trypsinized, pelleted, washed again with PBS, and snap-frozen in liquid nitrogen. Total RNA was extracted using the NucleoSpin RNA Plus kit (Macherey–Nagel, 740984.250) according to the manufacturer’s instructions.

Libraries were prepared from 500 ng total RNA using the QuantSeq 3′ mRNA-Seq FWD kit (Lexogen), pooled, and sequenced on an Illumina NovaSeq 6000 platform as part of a 100 bp paired-end run, yielding ∼4-5 million reads per sample. Only Read 1 was used for downstream analysis, following Lexogen guidelines. Reads were quality-checked (Phred > 30), adapter-and poly(A)-trimmed, aligned to the human reference genome (GRCh38) using STAR (v2.7), and counted with HTSeq (v2.0.2). Genes with fewer than 10 counts in at least three samples were excluded prior to differential expression analysis. Differential expression analysis was performed in R using DESeq2 (v1.44.0) with default normalization and statistical testing, and log_2_ fold-changes were shrunk using the adaptive shrinkage (“ashr”) method. For visualization of transcriptional responses, normalized counts from DESeq2 were used to calculate log_2_ fold changes relative to DMSO controls at the replicate level. A heatmap was generated using a published set of core estrogen target genes (*11*). Gene set enrichment analysis (GSEA) was performed using ranked gene lists based on log_2_ fold changes and running enrichment scores for the Human MSigDB HALLMARK_ESTROGEN_RESPONSE_EARLY (*50*). Transcriptional response was calculated as average log_2_FC across 190 genes detected as expressed in MCF7 cells out of 200 genes contained in the Human MSigDB HALLMARK_ESTROGEN_RESPONSE_EARLY (M5906) gene set were visualized as summary line plots across conditions.

### RNA-sequencing

MCF7 wild-type cells were seeded in 6-well plates at 5 × 10^4^ cells per well (DMEM + 10% FBS). After 72 h of hormone deprivation (phenol red-free DMEM supplemented with 10% charcoal-stripped FBS) cells were pre-treated for 1 h with DMSO or 2 µM ML792, and then further treated for 24 h with DMSO, 10 nM Estradiol, 10 nM Fulvestrant, or 10 nM 4-OHT. Following treatment, cells were washed once with ice-cold PBS and lysed directly in-well with 350 µL lysis buffer. Total RNA was isolated using the NucleoSpin RNA Plus kit (Macherey-Nagel, 740984.250) according to the manufacturer’s instructions.

Libraries were prepared using a reverse/second-strand ssRNA-seq protocol and sequenced on an Illumina NovaSeq X Plus platform (paired-end 150 bp). Reads were quality-checked, adapter- and poly(A)-trimmed, aligned to the human reference genome (GRCh38) using STAR (v2.7), and assigned to genes with featureCounts (v2.0.1). Genes with <10 counts in at least three samples were excluded prior to differential expression analysis. Differential expression analysis was performed in R (v4.4.1) using DESeq2 (v1.44.0) with default normalization and statistical testing, and log_2_ fold changes were shrunk using the adaptive shrinkage (“ashr”) method. For visualization of transcriptional responses, normalized counts from DESeq2 were used to calculate log_2_ fold changes relative to DMSO controls at the replicate level. A heatmap was generated using a selected gene set of core estrogen target (*11*) genes. GSEA was performed using ranked gene lists based on log_2_ fold changes, and running enrichment scores for the Human MSigDB HALLMARK_ESTROGEN_RESPONSE_EARLY (M5906) gene set (*50*) were visualized as summary line plots across conditions.

### CellTiter-Glo cell viability assay

MCF7 cells were seeded in 96-well plates at a density of 250 cells per well in 50 µL DMEM per well. The following day, 50 µL of 2x compound solutions in full growth medium were added for a final volume of 100 µL. Cells were grown in the presence of compounds for 7 days in a humidified incubator at 37⁰C and 5% CO_2_. 100 µL CellTiter-Glo reagent (G7570, Promega) per well was added, the plates were left to shake for 15 minutes, before measuring luminescence using a VICTOR X3 multilabel plate reader (Perkin Elmer) operated on PerkinElmer 2030 software (v4.0) or a GloMax Discover plate reader (Promega). Luminescence signal was normalized to DMSO-treated wells per plate and 4-parameter log-logistic curves were fitted in R. AOC (area over the growth curve) values were derived and normalized to obtain percent growth inhibition (fig. S7I), with 100% growth inhibition corresponding to 0% viability at all tested compound concentrations and 0% growth inhibition corresponding to no drug effects compared to DMSO at all tested doses.

### Competitive growth assay

MCF7 iCas9 cells were transduced with lentivirus expressing either an individual sgRNA or an array of three sgRNAs, either coupled to GFP and a blasticidin resistance gene (AAVS1, 3xCTRL) or mCherry and puromycin resistance gene (all test sgRNAs). After antibiotic selection, mCherry and GFP labelled cells were mixed in a 1:1 ratio. The number of expressed sgRNAs was matched between GFP-labelled control and mCherry-labelled test cell lines. The exact ratios of GFP and mCherry labelled cells in each pool were quantified by flow cytometry before and in regular intervals after doxycycline-based Cas9 induction (1 µg/mL). Three days after Cas9 induction, drug treatments were started, and compounds were replenished after every split and FACS analysis for at least 14 days. For analysis, absolute percentage of mCherry positive cells in each pool were normalized to start of treatment (day 0), relative changes between individual timepoints were linearly interpolated, values were averaged over replicates and plotted against pseudotime as mean ± SEM.

### Crystal violet colony formation assay

MCF7 iCas9 cells were transduced with lentivirus expressing AAVS1, CRBN, 3xCTRL or 3xRNF targeting sgRNAs and selected for sgRNA expression. Cas9 expression was induced with doxycycline (1 µg ml−1) for three days, before 2000 cells were seeded per well in 6 well plates. The following day, cells were treated with DMSO, dBET6 (250nM), SERDs (0.5 nM Fulvestrant or AZD9496, 0.25 nM Giredestrant or Camizestrant) or 4-OHT (2.5 nM). Cells were grown for 13 days under continuous treatment. For analysis, cells were washed with cold PBS, fixed with 1% PFA on ice for 30 minutes, washed with PBS, stained with Crystal Violet solution (0.1% in 10% EtOH) for 15 minutes at RT while shaking before washing with PBS three times. Plates were air-dried before imaging with a Nikon D700 DSLR camera. Images were processed with a linear gradient exposure mask and ‘Match Total Exposures’ function in Adobe Lightroom Classic (v14.0.1). Processed images were analyzed using a custom Python script in Spyder IDE (v6.0.7). For each plate, wells were detected via Hough circle transform and two background regions outside the wells were defined for threshold calibration. Crystal violet stained cells in each well were identified as pixels surpassing (i) a violet hue threshold (ii) a background-normalized saturation threshold and (iii) an average red/blue minus green pixel intensity threshold. Colonies were quantified as percent of stained pixels per well.

### FACS-based CRISPR-Cas9 ERα stability screens

The FACS-based CRISPR/Cas9 ERα stability reporter screens, library preparation, next-generation sequencing, and data analysis were performed as previously described (*81*). Briefly, MCF7 cells harboring a doxycycline-inducible Cas9 allele (iCas9) and expressing an ERα-GFP-P2A-mCherry reporter were transduced with a ubiquitin-proteasome system (UPS)-focused library (*78*), and KBM7 iCas9 (*79*) cells expressing an ERα-BFP-P2A-mCherry reporter were transduced with a genome-wide sgRNA library (*79*), both at multiplicity of infection of 0.1 and 1,000× library representation, respectively. Following selection for sgRNA library expression with G418 (1 mg/mL; Sigma-Aldrich, A1720), Cas9 expression was induced with doxycycline (0.4 µg ml−1; PanReac AppliChem, A2951) for three days, before cells were treated with DMSO or SERDs (MCF7: Fulvestrant, Giredestrant or Camizestrant, 10 nM for 6 h each; KBM7: Fulvestrant or Giredestrant, 100 nM each, or AZD9496 500 nM for 16 h each). Screens were performed in two biological replicates, except for DMSO screens in MCF7 cells, which were performed in three independent clones. Cells were harvested by trypsinization (MCF7 only), incubated for 10 min at 4 °C with anti-Thy1.1-APC antibody (BioLegend, 202526, 1:400) and Human TruStain FcX Fc receptor blocking solution (BioLegend, 422301, 1:400), fixed with 4% BD Fixation Buffer (BD Biosciences, 554655) for 45 min at 4 °C, protected from light, and stored overnight at 4 °C in PBS supplemented with 5% FBS and 1 mM EDTA. The following day, cells were sorted on a BD FACSAria Fusion (BD Biosciences) using a 70 µm nozzle. Aggregates, debris, Cas9-negative, and sgRNA library-negative (Thy1.1-APC-negative) cells were excluded. Remaining cells were sorted based on ERα-BFP or ERα-GFP and mCherry fluorescence into ERα^high^ (∼5%), ERα^mid^ (∼30%), and ERα^low^ (∼5%) populations, maintaining a minimum library coverage of 500× (genome-wide) or 1,000× (UPS-focused) per replicate.

Genomic DNA from sorted cell populations was isolated by cell lysis (10 mM Tris-HCl, 150 mM NaCl, 10 mM EDTA, 0.1% SDS), followed by proteinase K digestion (NEB #P8107) and DNase-free RNase treatment (Roche, 11119915001). DNA was purified by two rounds of phenol extraction (Sigma-Aldrich, P4557) and isopropanol precipitation (Sigma-Aldrich, I9516). Lentivirally integrated sgRNA expression cassettes were amplified by a two-step PCR using AmpliTaq Gold polymerase (Thermo Fisher Scientific, 4311818), introducing sample-specific barcodes in the first PCR and Illumina adapters in the second PCR. PCR products were purified using Mag-Bind TotalPure NGS beads (Omega Bio-tek, M1378-01), pooled, and sequenced on an Illumina NovaSeq 6000 platform.

Sequencing data were processed using a publicly available crispr-process-nf Nextflow pipeline (https://github.com/ZuberLab/crispr-process-nf/). Briefly, raw FASTQ files were trimmed with cutadapt (v4.4) to remove random barcodes and spacer sequences and demultiplexed based on sample barcodes. Reads were aligned to the sgRNA library using Bowtie2 (v2.4.5), and sgRNA abundance was quantified with featureCounts (v2.0.1). Gene-level enrichment analysis was performed using the crispr-mageck-nf workflow (https://github.com/ZuberLab/crispr-mageck-nf/). For combined analysis of MCF7 SERD screens (Fig. 1C), individual screens performed with Fulvestrant, Camizestrant and Giredestrant (fig. S1D-F) were combined as replicates in MAGeCK. Gene-level enrichment score was calculated as -log_10_ (P value) x log_2_ (fold-change).

### Viability-based CRISPR-Cas9 screens

For viability-based CRISPR-Cas9 screens, MCF7 iCas9 cells were transduced with a genome-wide sgRNA library and selected for library expression as above. Library expressing cells were treated with dox to induce Cas9 expression and after three days, cells were treated with DMSO or Fulvestrant (10 nM). Cells were washed, trypsinized and reseeded every three to four days. Treatment was renewed each time. After 14 days of treatment, surviving cells were harvested. At endpoint a large fraction of Fulvestrant treated cells had formed mammosphere-like spheroidal structures that were harvested and processed separately. Genomic DNA isolation and NGS library preparation was performed as described above for the FACS-based CRISPR/Cas9 screens. For Fulvestrant screens, adherent cell samples were lost during library preparation and only mammosphere-like samples were processed. Bioinformatic analysis of sequencing data was performed as described above. Gene-level enrichment was calculated in MAGeCK (0.5.9) (*84*) by comparing Fulvestrant-treated (spheroidal) populations to the DMSO-treated reference populations, using median-normalized read counts and replicate-level variance estimation. Gene-level enrichment score was calculated as -log_10_ (P value) x log_2_ (fold-change).

### Flow-cytometric ERα protein stability assay

To generate a stable ERα protein stability reporter cell line, MCF7 iCas9 cells were transduced with lentivirus (pLenti-SFFV-ERα-GFP-P2A-mCherry) expressing an ERα-GFP fusion coupled to an mCherry normalizer via a P2A ribosomal skipping element. For screen validation and to quantify the influence of genetic perturbations on SERD-based ERα degradation, reporter cells were transduced with lentiviral vectors expressing individual sgRNAs (pLenti-U6-sgRNA-IT-EF1αs-THY1.1-P2A-NeoR), or combinations of two or three sgRNAs (pLentiDual-hU6-sgRNA-IT-mU6-sgRNA-IT-EF1αs-THY1.1-P2A-NeoR or pLentiTriple-bU6-sgRNA-IT-hU6-sgRNA-IT-mU6-sgRNA-IT-EF1αs-mCherry-P2A-PuroR, respectively) to around 20-70% transduction efficiency. Cas9 expression was induced with doxycycline (1 µg ml−1) for 5-6 days, followed by 6 h of SERD treatment. For THY1.1 expressing sgRNA vectors, cells were stained with an APC-conjugated anti-mouse Thy1.1 antibody (202526, BioLegend; 1:400) and human TruStain FcX Fc receptor blocking solution (422302, BioLegend; 1:400) for 5 min in FACS buffer (PBS containing 5% FBS and 1 mM EDTA) at 4 °C. Cells were washed and resuspended in FACS buffer and analysed on an LSR Fortessa (BD Biosciences) operated on BD FACSDiva software (v9.0).

Flow-cytometric data analysis was performed in FlowJo v10.10.1 (Becton Dickinson). ERα-GFP and mCherry mean fluorescence intensity values for each sample were normalized by background subtraction of the respective values from reporter-negative MCF7 cells. ERα abundance was calculated as the ratio of background subtracted ERα-GFP to mCherry mean fluorescence intensity, and is displayed normalized to DMSO-treated sgRNA negative cells. For statistical analysis, two-way ANOVA with Tukey HSD post-hoc test was performed in R Studio.

### Immunoblotting

Cells were washed with ice-cold PBS supplemented with 10 mM N-ethylmaleimide (NEM; Thermo Fisher Scientific, #11891335) after treatment and harvested by scraping. Cell pellets were lysed in RIPA buffer containing 0.6% SDS, supplemented with benzonase (1:1000; Sigma, 70746), Halt protease inhibitor cocktail (1:100; Thermo Fisher Scientific, 11804111), and 20 mM NEM for 30 min on ice. After three rounds of sonication (30 seconds each, either in a Diagenode Bioruptor Pico or a QSonica Q800R3), lysates were cleared by centrifugation at maximum speed for 15 min at 4 °C, and protein concentrations were determined using Pierce BCA protein assay (Thermo Fisher Scientific, PI23225). Equal amounts of protein (20-30 µg) were mixed with 4× Bolt LDS sample buffer (Thermo Fisher Scientific, B0008) and 50 mM dithiothreitol (DTT), denatured at 95 °C for 5 min, and resolved on Bolt 4–12% Bis-Tris Plus WedgeWell gels (Thermo Fisher Scientific). Proteins were transferred to nitrocellulose membranes (Cytiva, 10600002), blocked for 1 h at room temperature in 5% non-fat dry milk in Tris-buffered saline containing 0.1% Tween-20 (TBS-T), and incubated with primary antibodies overnight at 4 °C. Membranes were then incubated with horseradish peroxidase (HRP)–conjugated secondary antibodies for 1 h at room temperature and imaged using a ChemiDoc MP Imaging System (Bio-Rad) with ECL substrate (Bio-Rad, 1705060). Primary antibodies used were β-actin (1:10,000, Sigma-Aldrich, A5441), ERα (1:1,000, Cell Signaling Technology, #13258), ERα (1:2000, Abcam, ab16460), ERα (1:2000, MedChemExpress, HY-P80663), GAPDH (1:2,000, Santa Cruz Biotechnology, sc-47724), RNF4 (1:1,000, a gift from Ronald Hay), RNF111 (1:1,000, Novus Biologicals, H00054778-M05), RNF216 (Biomol, #A304-111A), FLAG-Tag (Sigma, F1804), PIAS1 (Cell Signaling Technology, #3550), PIAS4 (Cell Signaling Technology, #4392), CBP (Cell Signaling Technology, 7389), NCOA3 (Cell Signaling Technology, #2126)m, SUMO2/3 (Abcam, ab81371). Secondary antibodies used were anti-rabbit IgG-HRP (Cell Signaling Technology, #7074, 1:10,000), anti-mouse IgG–HRP (Cell Signaling Technology, 7076, 1:10,000), anti-chicken IgY–HRP (Thermo Fisher Scientific, A16054), anti-mouse IgG-IRDye 800cw (1:10000, LI-COR Bio, 926-32210), anti-rabbit IgG-IRDye 800cw (1:10000, LI-COR Bio, 926-32211).

### Immunoprecipitation

For immunoprecipitation of ERα from MCF7 cells, sgRNA expressing iCas9 cells were treated with doxycycline for 7 days to induce 3xCTRL or 3xRNF knockout. Cells were then pre-treated with Carfilzomib (1 μM; MedChem Express, #HY-10455) for 1 h in absence or presence of the SUMOi ML792 (2 μM; MedChem Express, #HY-108702), followed by treatment with Fulvestrant (10 nM) or vehicle (DMSO) for 6 h. Cells were washed with ice-cold PBS supplemented with 10 mM N-ethylmaleimide (NEM; Thermo Fisher Scientific, #11891335) harvested by scraping. Cell pellets were lysed in RIPA buffer containing 0.6% SDS, supplemented with benzonase (1:1000; Sigma, 70746), Halt protease inhibitor cocktail (1:100; Thermo Fisher Scientific, 11804111), and 20 mM NEM for 30 min on ice. Following three rounds of sonication (30 seconds each, using a QSonica Q800R3), lysates were cleared by centrifugation at maximum speed for 15 min at 4 °C, and protein concentrations were determined using Pierce BCA protein assay (Thermo Fisher Scientific, PI23225). Before immunoprecipitation, per sample 7.5 μL Invitrogen Protein A Dynabeads (Fisher Scientific, # 10334693) were washed, incubated with 2 μL α-ERα antibody (Cell Signaling Technology, #8644) or concentration-matched IgG Isotype Control (Cell Signaling Technology, #3900) for 4 hours while rotating at 4 °C). After excess antibody was removed by washing with RIPA buffer supplemented with 0.5% BSA (Sigma-Aldrich, #A9647), beads were incubated with 500 μg protein lysates over night while rotating at 4 °C. The following day, beads were washed and protein was eluted by boiling beads in 4x LDS sample buffer 4× Bolt LDS sample buffer (Thermo Fisher Scientific, B0008) in the presence of 50 mM dithiothreitol (DTT) for 15 min. Immunoprecipitated protein was analyzed via immunoblotting as described above. Secondary antibody used for immunoblotting was HRP-conjugated anti-rabbit IgG light-chain specific (Cell Signaling Technology, #93702).

### Split-HaloTag interaction assay

HEK293T cells co-transduced with EGFP-cpHaloΔ-ERα and Hpep3-SUMO3-EF1α-TagBFP were seeded in 96-well plates at a density of 6 × 10^4^ cells per well. The following day, cells were pre-treated with Carfilzomib (1 µM; MedChem Express, #HY-10455) for 1 h in absence or presence of the SUMOi ML792 (2 µM; MedChem Express, #HY-108702), followed by compound treatment in combination with HaloTag-TAMRA ligand (100 nM; MedChemExpress, HY-D2270) for 6 h. To harvest samples, cells were trypsinized, centrifuged and resuspend in FACS buffer (PBS containing 5% FBS and 1 mM EDTA) before acquiring on an LSR Fortessa (BD Biosciences) operated on BD FACSDiva software (v9.0).

### NanoBiT ERα interaction assay

HEK293 cells were seeded in white 96-well plates at a density of 5 × 10⁴ cells per well. The following day, cells were co-transfected using Lipofectamine 3000 (Thermo Fisher Scientific, #15292465) with 50 ng ERα-LgBiT and 50 ng of the indicated SmBiT fusion construct (PIAS1, PIAS4, MTA2 or HDAC1; 100 ng total DNA per well). 48 h later, cells were pre-treated with Carfilzomib (1 µM; MedChem Express, #HY-10455) for 1 h in the absence or presence of the SUMOi ML792 (2 µM; MedChem Express, #HY-108702), followed by addition of the Nano-Glo Endurazine live-cell substrate (Promega, #2571). Baseline luminescence (t = 0) was recorded, after which cells were treated with Estradiol or Fulvestrant (10 nM each) for the indicated timespans. Bioluminescence, reporting on the reconstitution of the split NanoLuc upon ERα-prey proximity, was measured on a GloMax Discover plate reader (Promega).

### Protein expression and purification

STREP-MBP-TEV-ERα full-length (fl) (UniProt entry: P03372, residues 1–595), STREP-MBP-TEV-ERα^DBD-LBD^ (Residues 174–595), FLAG-SUMO2(1–92)-SUMO2(1–92)-TEV-ERα^fl^, STREP-Halo-TEV-RNF111 (UniProt entry: Q6ZNA4, residues 221–994), FLAG-MBP-TEV- PIAS1^fl^ (UniProt entry: O75925), FLAG-TEV-PIAS4^fl^ (UniProt entry: Q8N2W9), FLAG-TEV-PIAS4^ΔSAP^ (Residues 118-510), FLAG-TEV-PIAS4^ΔSAP,PINIT^ (Residues 279-510) and the *de novo* designed His-TEV-ERα^LBD^ binder were cloned into a pAC8-derived vector and recombinantly expressed in *Trichoplusia ni* High Five^TM^ (BTI-Tn-5B1-4) insect cells (Thermo Fisher, B85502) using the flashBAC^TM^ baculovirus expression system (*85*). The cells were cultured at 27 °C in SF4 Baculo Express Media (BioConcept, 9-00F38-K) and harvested 40 h after induction of protein expression. For FLAG-SUMO2-SUMO2-TEV-ERα^fl^, 10 µM of Estradiol or Fulvestrant was added to the cultures 16 h post transduction, for STREP-MBP-TEV-ERα^fl^, 1 µM of Estradiol was added to facilitate protein expression. To obtain Ful-bound STREP-MBP-TEV-ERα^fl^, Estradiol added during protein expression was displaced by Ful supplied in the purification buffers (see below).

Briefly, protein expressing cells were harvested and lysed by sonication in lysis buffer (50 mM HEPES pH 8.0, 150 mM KCl, 10 mM MgCl_2_, 0.5 mM CaCl_2_, 10% glycerol, 0.5 mM TCEP (for STREP-tagged proteins), supplemented with 0.05 mg/ml DNase (Sigma-Aldrich, DN25) and 1x protease inhibitor cocktail (MedChemExpress, HY-K0010)). For STREP- and FLAG-tagged ERα^fl^, 10 µM E2 or Ful was added to the lysis buffer, whereas STREP-tagged ERα^DBD-LBD^ was initially purified in the absence of ligand. To enhance ligand solubility in aqueous solution, 0.2% (w/v) 2-Hydroxypropyl-β-cyclodextrin (Sigma-Aldrich, H107-5G) was added to the buffer per 10 µM final ligand concentration. For STREP-tagged ERα constructs 4 mM CHAPS, for FLAG-tagged SUMO2-SUMO2-ERα^fl^, PIAS1^fl^, and PIAS4 constructs, 0.1% Triton X-100 was added to the lysis buffer. Cells expressing STREP-tagged RNF111 were lysed in lysis buffer containing 300 mM KCl and 0.1% Triton X-100. Subsequently, the lysates were cleared by ultracentrifugation (40000 rpm, 45 min, 4 °C) and the filtered supernatant was used for affinity purification.

The supernatant containing FLAG-tagged SUMO2-SUMO2-ERα^fl^, PIAS1^fl^ or PIAS4 constructs was incubated with FLAG M2 gel (Sigma-Aldrich, A2220) for 90 min at 4 °C. After sequential washes with wash buffer (50 mM HEPES pH 8.0, 10% glycerol, 0.1% Tween-20) supplemented with 150 mM KCl, 700 mM KCl, and 150 mM KCl with 5 mM MgCl_2_ and 4 mM ATP, elution was performed with wash buffer containing 150 mM KCl, and 0.4 mg/ml 3xFLAG peptide (Sigma-Aldrich, F4799). For SUMO2-SUMO2-ERα^fl^, 10 µM of Estradiol or Fulvestrant was added to the wash buffers. The protein-containing fractions were pooled, concentrated, and purified by size exclusion chromatography (SEC) on a Superdex 200 (S200) column (Cytiva, 28990944) equilibrated in SEC buffer (25 mM HEPES pH 8.0, 5% glycerol, 0.5 mM TCEP) supplemented with 300 mM NaCl for SUMO2-SUMO2-ERα^fl^ or 150 mM NaCl for PIAS1/4 constructs. FLAG-TEV-PIAS4^ΔSAP,PINIT^ was applied onto a Superdex 75 (S75) column (Cytiva, 28989333), diluted to 100 mM and finally purified via a MonoQ column (Cytiva, 17516601) upon elution in a linear gradient from 150 mM to 1 M NaCl with fractions pooled and subsequently concentrated.

The supernatant containing STREP-tagged ERα constructs and RNF111 was incubated with StrepTactin (IBA Lifesciences, 2-1201-025) resin for 60 min at 4 °C. After sequential washes with wash buffer (50 mM HEPES pH 8.0, 10% glycerol, 0.5 mM TCEP) supplemented with 150 mM KCl, 700 mM KCl, and 150 mM KCl with 5 mM MgCl_2_ and 4 mM ATP, elution was performed with wash buffer containing 150 mM KCl and 5 mM Desthiobiotin (IBA lifesciences, 2-1000-005). For the ERα constructs, wash buffer was always supplemented with 0.5 mM CHAPS and for ERα^fl^ additionally contained 10 µM of either Estradiol or Fulvestrant. For RNF111, wash buffer included 0.1% Triton X-100. Elution fractions containing RNF111 were pooled, concentrated, and directly subjected to a Superose 6 (S6) column (Cytiva, 29091596) equilibrated in SEC buffer (25 mM HEPES pH 8.0, 300 mM NaCl, 10% glycerol, 0.5 mM TCEP).

The STREP elution fractions of the ERα constructs were directly applied to a Heparin column (Cytiva, 17040703) equilibrated in wash buffer supplemented with 150 mM NaCl, 0.2 mM CHAPS and 2 µM of either Estradiol or Fulvestrant for ERα^fl^. The proteins were eluted in a linear gradient from 150 mM to 1 M NaCl. Elution fractions were pooled, the salt concentration was lowered to 100 mM NaCl and applied onto a Mono Q column (Cytiva, 17516601) for ERα^fl^ or HiTrap^TM^ Q (Cytiva, 17115401) for apo ERα^DBD-LBD^ equilibrated in wash buffer with 100 mM NaCl, 0.2 mM CHAPS and 2 µM of Estradiol or Fulvestrant for ERα^fl^. The proteins were eluted in a linear gradient from 100 mM to 1 M NaCl with fractions pooled and subsequently concentrated. ERα^fl^ and apo ERα^DBD-LBD^ were directly applied onto a S6 column (Cytiva, 29091596) equilibrated in SEC buffer (25 mM HEPES pH 8.0, 5% glycerol, 0.5 mM TCEP) supplemented with 300 mM KCl, 0.5 mM CHAPS, 10 µM Estradiol or Fulvestrant for ERα^fl^ or 150 mM NaCl, 0.2 mM CHAPS for apo ERα^DBD-LBD^. To obtain ligand-bound ERα^DBD-LBD^, the apo protein was dialyzed overnight against SEC buffer including excess Estradiol or Fulvestrant and subsequently purified by a S6 column under identical conditions.

Ubiquitin (UniProt entry: P0CG47, residues 1–76, with N-terminal SGSCG linker), SUMO1 (UniProt entry: P63165, residues 2–97, S9C, C52A), SUMO2 (UniProt entry: P61956, residues 2–93, A2C, C48A), SUMO3 (UniProt entry: P55854, residues 2–92, A2C, C47A), Ubc9^fl^ (UniProt entry: P63279) were cloned into bacterial pET-28a vectors and expressed as N-terminal His_5_- (ubiquitin) or His_10_-TEV-tagged proteins. RNF4^fl^ (UniProt entry: P78317), PIAS4^PINIT-SP-RING^ (Residues 118-416), ERα^LBD^ (Residues 301–552) or BRIL-ERα^LBD^ (Residues 312–552^Y331F, D332S, L462T, Δ333-338, Δ463-465^) were cloned into bacterial pET-28a vectors and expressed as N-terminal His_10_-MBP-TEV (RNF4^fl^) or His_10_-SUMOstar-TEV tagged proteins, respectively. All recombinant proteins were produced in *E. coli* BL21-CodonPlus (DE3)-RIL cells (Agilent, 230245). Cells were grown at 37 °C in LB media. At OD_600_ 0.5–0.7 protein expression was induced by the addition of IPTG. For SUMO1-3 and Ubc9^fl^ 0.5 mM IPTG was added following expression at 37 °C for 4 h or 30 °C for 5 h, respectively. For ubiquitin, 1 mM IPTG was added following expression at 30 °C for 6 h. For the ERα^LBD^ constructs and PIAS4^PINIT-SP-RING^, protein expression was induced with 0.3 mM IPTG following expression at 16 °C for 18 h. RNF4 was expressed as described previously (*86*).

Cell pellets from protein expression in *E. coli* or from insect cells containing the His-TEV-ERα^LBD^ binder were resuspended in lysis buffer (50 mM HEPES pH 8.0, 500 mM NaCl, 5% glycerol, 0.5 mM TCEP, 10 mM MgCl_2_, 0.5 mM CaCl_2_ supplemented with 0.05 mg/ml DNase, 1x protease inhibitor cocktail, and 20-40 mM imidazole), lysed by sonication, and clarified by ultracentrifugation. The supernatant was applied to a HisTrap^TM^ column (Cytiva, 17524802). After washing with wash buffer (50 mM HEPES pH 8.0, 500 mM NaCl, 5% glycerol, 0.5 mM TCEP, and 75 mM imidazole), proteins were eluted with wash buffer containing 500 mM imidazole.

For maleimide labelling of ubiquitin, SUMO2 and SUMO3, 5 mM TCEP was added and incubated for 45 min on ice. The proteins were buffer exchanged into PBS using a PD-10 desalting column (Cytiva, 17085101). Subsequently, 0.8 µmol of each protein was mixed with 2.4 µmol of IRDye 680RD Maleimide (LI-COR Bio, 929-71051) and incubated for 3 h at room temperature. Excess dye was quenched with 5 mM cysteine and 2.5 mM TCEP, and the labelled proteins were subsequently re-applied to a HisTrap^TM^ column equilibrated in wash buffer with 40 mM imidazole to remove unreacted dye. Labelled proteins were eluted in wash buffer containing 500 mM imidazole.

To cleave the His tag, the proteins were incubated with Tobacco Etch Virus (TEV, homemade) protease at a 1:30 (w/w) ratio and dialyzed overnight into wash buffer supplemented with 30 mM imidazole. Dialyzed samples were again loaded onto a HisTrap^TM^ column to remove TEV protease and His tag. The proteins were subsequently concentrated and purified by SEC equilibrated in SEC buffer (25 mM HEPES pH 8.0, 150 mM NaCl, 5% glycerol, 0.5 mM TCEP). SUMO2/3 either His-tagged or cleaved and labelled, as well as cleaved Ubc9, RNF4, and PIAS4^PINIT-SP-RING^ were applied onto a S200 column (Cytiva, 28989335, 28990944). Cleaved and labelled ubiquitin, cleaved ERα^LBD^ binder and non-cleaved His-SUMO1 were purified by a Superdex 75 (S75) column (Cytiva, 28989333). Cleaved apo ERα^LBD^ was purified directly using an S75 column (Cytiva, 29148721). Ligand-bound ERα^LBD^ was generated by overnight dialysis against buffer including excess Estradiol or Fulvestrant, followed by purification under identical conditions in the presence of the respective ligand. Cleaved apo BRIL-ERα^LBD^ was purified using an S200 column and subsequently dialyzed overnight in the presence of Fulvestrant or Giredestrant.

Methylated SUMO2 was prepared from 220 nmol of His-tagged SUMO2 purified by S200 using the reductive alkylation kit (Hampton Research, HR2-434) according to the manufacturer’s instructions. After reductive methylation, SUMO2 was purified by S75 in SEC buffer.

#### *In vitro* SUMOylation assays

SUMO conjugation assays were conducted in 20 mM HEPES pH 7.4, 50 mM NaCl, 5 mM MgCl_2_, 1 mM DTT, 0.2% (w/v) 2-Hydroxypropyl-β-cyclodextrin, +/- 10 µM ligand or DMSO in a 20 µL reaction volume containing 50 nM SUMO E1 enzyme (Enzo, BML-UW9330), 100 nM Ubc9^fl^, 300 nM FLAG-tagged PIAS1^fl^,PIAS4^fl^ or truncated constructs or without SUMO E3 ligase, 5 µM of IRDye 680 maleimide-labelled SUMO2 and SUMO3 (fig. S3N, fig. S6B and I-L) or His-tagged SUMO1,2,3 and methylated SUMO2 (fig. 2E and fig. S3L) and 400 nM of the respective ERα substrate. SUMO reactions were initiated by the addition of 5 mM ATP and incubated at 30 °C for 20 min. SUMOylation assays in the presence DNA (fig. S6K) were performed with 40 nM of an oligonucleotide containing the estrogen response element (ERE) fused to a PIAS binding site (*87*) (Table S1) and incubated for 1 h. For SUMOylation assays in the presence of increasing ERα^LBD^ concentrations (fig. S6L), ERα^LBD^ pre-bound to Estradiol or Fulvestrant was titrated in twofold increments starting from 1:1 to 1:16 molar ratio (ERα^fl^ : ERα^LBD^). For SUMO assays in the presence of ERα^LBD^ binder (fig. S6B), the binder was titrated in twofold increments starting from 1:1 to 1:8 molar ratio (ERα^fl^ : ERα^LBD^ binder) For SUMO assays with various ligands (fig. S3N), 5 µM apo STREP-MBP-TEV-ERα^DBD-LBD^ was pre-incubated overnight with a 10-fold molar excess of the respective compound and used as stock for the SUMO assays. The reactions were stopped by adding 6 µL 4xSDS sample buffer (*88*) supplemented with 0.1 mM EDTA pH 8.0. 5 µL of each reaction was applied to either 3–8% Tris-acetate or 4–12% Bis Tris Plus gels (Invitrogen, TA03815BOX, NW04127BOX) and run at 150 V for 1 hr. After gel electrophoresis, the gel was transferred to a nitrocellulose membrane (Bio-Rad, 1704158) using the Trans-Blot Turbo Transfer System (Bio-Rad). Following blocking with 5% skim milk solution in TBS-T buffer, the membrane was incubated with an Anti-ERα antibody at 4 °C overnight (Blots including ERα^fl^: MedChemExpress, HY-P80663; blots including ERα deletions: Abcam, ab16460, both at 1:2000). The following day, the membrane was washed thrice with TBS-T and subsequently incubated with a secondary antibody conjugated to IRDye 800CW (LI-COR Bioscience, 926-32210, 926-32211, 1:10000) for 1 h at room temperature. After additional three washes with TBS-T, the membrane was imaged on a LI-COR Odyssey DLx imaging system (LI-COR, Biosciences) at 800 nm and 700 nm in the presence of labelled SUMO2/3.

For correlation analysis (fig. 2F), ERα immunoblots of *in vitro* SUMOylation reactions (fig. S3N) and cellular SUMOylation of endogenous ERα in MCF7 cells co-treated with Carfilzomib and ERα ligands (fig. S3M) were quantified via Bio-Rad Image Lab software (Version 6.1.0 build7). For each compound, ERα-SUMO signal intensity was normalized to non-SUMOylated ERα intensity and the log_2_ fold increase in ERα SUMOylation was calculated compared to DMSO. log_10_-transformed fold-changes over DMSO were plotted against fold-increase over DMSO of ERα-GFP retained in cell nuclei after sample processing according to the chromatin FACS protocol described above. Linear regression, Pearson R, and P value (Pearson’s two-sided correlation test) were calculated in R.

#### *In vitro* ubiquitylation assays

Ubiquitylation reactions shown in Fig. 2B were performed at 37 °C for 40 min in 20 mM HEPES pH 7.4, 50 mM NaCl, 5 mM MgCl_2_, 1 mM DTT, 0.2% (w/v) 2-Hydroxypropyl-β-cyclodextrin, +/-10 µM ligand or DMSO in a 20 µL reaction volume. The reactions contained 100 nM UBE1 E1 enzyme (R&D systems, E-305), 500 nM UbcH5a E2 enzyme (R&D systems, E2-616), 5 µM of IRDye 680 maleimide-labelled ubiquitin, 600 nM of the respective ERα^fl^ substrate and 500 nM of RNF4 or STREP-tagged RNF111. The reactions were initiated by the addition of 5 mM ATP and subsequently processed for immunoblotting as described in the section *in vitro* SUMOylation assays.

### **ER**α-PIAS4 *in vitro* binding assays

Binding assays with recombinant FLAG-PIAS4^fl^ and STREP-MBP-ERα^fl^ (fig. S3K) were performed using 8 µl of STREP-Tactin^®^ XT 4Flow^®^ slurry (IBA lifesciences, 2-5010-025) equilibrated in assay buffer (25 mM HEPES pH 8.0, 100 mM NaCl, 0.5 mM TCEP, 0.05% Tween-20 supplemented with 10 µM Estradiol (E2) or Fulvestrant in 0.2% (w/v) 2-Hydroxypropyl-β-cyclodextrin). After equilibration, excess buffer was removed and replaced with 100 µl of fresh assay buffer, followed by the addition of 10 pmol ERα^fl^. After incubation at 4°C for 90 min, the supernatant was removed, the beads were washed with 450 µl of assay buffer, followed by the addition of 100 µl of assay buffer together with 40 pmol of FLAG-PIAS4^fl^. The protein-bound resin was incubated for another 90 min followed by three sequential wash steps with 450 ul of assay buffer. The proteins were eluted by adding 20 µl of assay buffer with 6 µl of 5x SDS sample buffer (*88*). 12 ul of each sample was applied to a 4-12% SDS-PAGE gradient gel (Invitrogen, NW04127BOX) run at 150 V for 1 hr and subsequently processed for immunoblotting as described in the section *in vitro* SUMOylation assays using primary antibodies against ERα^fl^ (MedChemExpress, HY-P80663, 1:2000) and PIAS4 (Cell Signaling Technology, #4392, 1:1000) and an IRDye 800CW (LI-COR Bioscience, 926-32211, 1:10000) secondary antibody. Final imaging of the membrane was done using a LI-COR Odyssey DLx imaging system (LI-COR, Biosciences) at 800 nm.

### Fluorescence polarization (FP) assays

NCOA1 binding studies (fig. S3H) were performed using a fluorescein-labelled NCOA1 peptide (Residues: 686–698, Flc-ARHKILHRLLQEGS, UniProt: Q15788) as a fluorescent tracer (*22*). Increasing amounts of ERα^LBD^ either apo or pre-bound to Estradiol or Fulvestrant (0.6 nM–10 µM, final concentration) were mixed with labelled NCOA1 peptide (10 nM final concentration) in a 384-well microplate (Greiner Bio-One, 784076) at a final volume of 10 µL in buffer containing 25 mM HEPES pH 8.0, 75 mM NaCl, 2.5% glycerol, 0.0025% (v/v) Tween-20, 0.75 mM TCEP, 0.1% (w/v) 2-Hydroxypropyl-β-cyclodextrin, +/- 5 µM ligand or DMSO and incubated for 15 min at room temperature.

DNA binding studies (fig. S7G) were performed using a fluorescein-labelled Estrogen response element (ERE) DNA oligo (5’-Flc-CTCCAGGTCACCCTGACCTTAG-3’). STREP- MBP-TEV-ERα^DBD-LBD^ (Fixed final concentration of 0.5 µM) was pre-incubated with labelled ERE (Fixed final concentration of 10 nM) for 30 min on ice and used for back-titration with increasing amounts of the same unlabelled ERE (1.2 nM–5 µM, final concentration) at a final volume of 10 µL and buffer containing 25 mM HEPES pH 8.0, 75 mM NaCl, 2.5% glycerol, 0.1 mM CHAPS, 0.5 mM TCEP, 0.1% (w/v) 2-Hydroxypropyl-β-cyclodextrin, +/- 5 µM ligand or DMSO. The changes in FP were measured using a PHERAstar FS microplate reader (BMG Labtech) with excitation and emission wavelengths of 485 and 520 nm, respectively. Both experiments were performed in technical triplicates. The raw FP values were converted to fraction bound (F_bound_) according to:

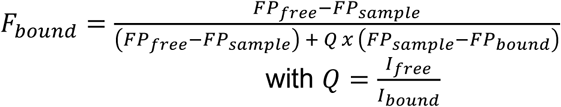

For FP experiments with NCOA1, FP_bound_/I_bound_ was defined as the highest FP value obtained upon binding of NCOA1 to 10 µM ERα^LBD^ in the presence of Estradiol. For FP experiments with ERE, FP_bound_/I_bound_ corresponded to ERα^DBD-LBD^ bound to labelled-ERE in the absence of unlabelled ERE. The fraction bound was plotted against increasing protein or DNA concentrations and fitted assuming a one-site binding model to determine the K_d_ or using an Inhibitor *vs* response – Variable slope model to calculate the IC_50_ using GraphPad Prism (Version 10.2.3).

### Design of ERα^LBD^ binders and BRIL-ERα^LBD^ for cryo-EM

BindCraft2 (manuscript in preparation) was utilized to design *de novo* binders (∼25–35 kDa) that structurally stabilize the ERα^LBD^ and increase its molecular mass. The dimeric ERα^LBD^ (PDB ID: 6ZOR), trimmed to residues 303–552, was used with the “largebinder” settings and without explicitly defining hotspots. The pipeline was run until 15 designs passed the default filters, and candidates were ranked by the i_pTM score predicted by AlphaFold2 (AF2) (*89*) to prioritize experimental testing. The top six candidates were co-expressed with full-length STREP-MBP-TEV-ERα^fl^ in *Trichoplusia ni* High Five^TM^ (BTI-Tn-5B1-4) insect cells (Thermo Fisher, B85502), and their binding ability was assessed by STREP pulldown of ERα^fl^. The best-performing binder was subsequently selected, expressed, and purified as described in the section Protein expression and purification and used for cryo-EM.

To generate the BRIL-ERα^LBD^ (Residues: 312–552^Y331F, D332S, L462T, Δ333–338, Δ463–465^), an 18-amino acid α-helical linker extending the N-terminal helix 1 from A312 was iteratively tested using AF3 (*90*) to enable in-frame fusion of the BRIL (*47*). The α-helical linker sequence was subsequently optimized with ProteinMPNN (*91*). The resulting engineered construct formed an ordered array of ERα^LBD^ dimers mediated by domain swapping of Helix 1 even in the absence of the ERα^LBD^ binder (data not shown). However, high-resolution structures of the Fulvestrant- and Giredestrant-bound states were obtained only in the presence of the binder, with two ERα^LBD^ dimers bound to a single binder molecule as the repeating unit (Fig. 2G-I, fig. S6A-F, fig. S4 and S5).

### Cryo-EM sample preparation and data collection

Purified BRIL-ERα^LBD^ bound to Fulvestrant and the ERα^LBD^ binder were mixed at ∼150 µM and ∼230 µM, respectively. Following incubation on ice for 30 min, the sample was applied onto a Superdex 200 Increase 3.2/300 (Cytiva, 28990946) equilibrated in cryo-EM buffer (25 mM HEPES pH 8.0, 100 mM NaCl, 0.5 mM TCEP and 10 µM Fulvestrant in 0.2% (w/v) 2-Hydroxypropyl-β-cyclodextrin). Peak fractions containing the complex were pooled and concentrated to ∼30 µL using an Amicon centrifugal filter concentrator (10 kDa MWCO, Merck-Millipore, UFC501096). To prepare the Giredestrant-bound BRIL-ERα^LBD^ complex, ∼100 µM of apo BRIL-ERα^LBD^ was mixed with ∼80 µM of ERα^LBD^ binder in the presence of ∼400 µM Giredestrant in cryo-EM buffer supplemented with 1% (w/v) 2-Hydroxypropyl-β-cyclodextrin and incubated overnight on ice. The sample was directly used to prepare cryo-EM grids. For grid preparation, 4 µL of each sample was applied to glow-discharged Quantifoil^TM^ R1.2/1.3 200 gold mesh grids (Quantifoil Micro Tools). After 5 sec, the sample was blotted for 5 sec and subsequently plunge-frozen into liquid ethane at 95% humidity and 4 °C using Vitrobot Mark IV (FEI).

For the Fulvestrant-bound complex, two datasets with 3663 and 17410 movies were collected on a Cs-corrected (CEOS GmbH, Heidelberg, Germany) Titan Krios electron microscope (Thermo Fisher Scientific) operating at 300 kV. For Giredestrant, 7670 movies were collected on a Glacios electron microscope (Thermo Fisher Scientific) operating at 200 kV. Automatic data collection was performed using the EPU software (Thermo Fisher Scientific) at a nominal magnification of 75,000x and 120000x corresponding to a pixel size of 0.845 Å for the Krios and 0.84 Å for the Glacios datasets, respectively. Movies were recorded in EER format with a Falcon 4i direct electron detector (Thermo Fisher) with a total electron dose of 50 e−/Å2. Defocus values ranged from -0.8 to -2.0 μm.

### Cryo-EM data processing and model building

On-the-fly postprocessing of raw data was performed with CryoFLARE v1.13 (*92*), which utilizes WarpTools (v2.0.0dev36; accessed via doi.org/10.5281/zenodo.13982246) for motion correction and CTF estimation. Movies were fractionated into 50 frames, corresponding to 1 e−/Å2 per frame. Data processing for all datasets was performed in cryoSPARCv4.7.1 (*93*), with additional particle picking in crYOLO (v1.9.9) (*94*) as detailed in fig. S4 and S5. For Fulvestrant, crYOLO-based particle picking of the smaller dataset, followed by *ab initio* reconstruction, and 3D refinement, produced an initial model based on 142246 particles at ∼4.1 Å resolution. This model was then used for template-based particle picking of the larger dataset. Combined with crYOLO-based picking and blob picker, this yielded ∼1.5 million particles, which underwent four rounds of heterogeneous refinement. These particles were then combined with the similarly classified particles from the small dataset for non-uniform refinement, resulting in a tetrameric map of two ERα^LBD^ dimers bound to one ERα^LBD^ binder (fig. S4). To resolve the dynamics at the compound binding sites, 3D variability analysis (3DVA) was applied with a mask covering the dimer resolved at higher resolution (dimer^1^). 3DVA revealed one binding site containing Fulvestrant tail density without helix 12 (H12) density (fig. 2G-I), whereas the second site retained residual H12 density (fig. S6C and D). This likely reflects the conformational flexibility of the Fulvestrant tail or mixed populations of Fulvestrant-bound and ligand-free states, which may permit H12 folding at this site despite its incompatibility based on static modelling. The same two binding modes were observed for the other dimer (dimer^2^), despite its overall lower resolution. Subsequent non-uniform refinement yielded a consensus map based on 130795 particles at 3.2 Å resolution, which was used to generate a composite cryo-EM density map based on individual local refinements for dimer^1^, dimer^2^ and binder (fig. S4).

For the Giredestrant dataset, particles were likewise picked using crYOLO-based particle picking. After particle extraction, one round of 2D classification and two rounds of heterogeneous refinement using *ab initio* volumes derived from the small Fulvestrant dataset, resulted in 328261 particles. The re-extracted non-binned particles were subsequently used for final non-uniform refinement to obtain a cryo-EM map based on 319580 particles at 3.5 Å resolution. In contrast to the Fulvestrant-bound complex, all Giredestrant binding sites show comparable H12 densities.

Reported resolution values for the final maps after non-uniform refinement are based on the gold-standard Fourier shell correlation (FSC) curve at 0.143 criterion (*95*). High-resolution noise substitution was used for correcting the effects of soft masking for the related FSC curve. Local resolution was estimated in cryoSPARCv4.7.1.

For model building of the Fulvestrant-bound complex, a composite cryo-EM map was first generated in ChimeraX (v1.10.1) using the volume maximum command (*96*) based on the final local refinement maps for dimer^1^, dimer^2^, and binder (fig. S4). The Giredestrant-bound complex was modelled using the map obtained, after the final non-uniform refinement (fig. S5). Models for the BRIL-ERα^LBD^ fusion construct in complex with ERα^LBD^ binder were predicted with AlphaFold3 (AF3) (*90*) and fitted into the maps using ChimeraX (v1.10.1) (*96*) fit-in-map. Model adjustments were done in COOT (v0.9.8.1) (*97*). Fulvestrant and Giredestrant were placed using the PDB chemical ID FVS or ZNM, respectively, as a template. Helix 1, involved in domain swapping between protomers and involved in tetramer formation was modelled. The models were refined by flexible fitting in ISOLDE (*98*) using self- and secondary structure restraints, and side chains were adjusted manually in COOT and ISOLDE (v1.8). For the protomer showing overlapping density between Fulvestrant and helix 12 (H12) (fig. S5), the occupancy was set to 0.5 with the ligand assigned to alternative conformation A and H12 to alternative conformation B. Finally, the models were refined in PHENIX (*99*) using coordinate restraints with a sigma of 0.5. B-factors and occupancies were refined in the same step. Validation was carried out in PHENIX (v1.20.1), including MOLPROBITY (*100*) and EMRinger (*100*). Figures were prepared with ChimeraX (v1.10.1).

### Orthotopic xenograft model

The xenograft studies were approved by the Austrian Ministry (approval number MA58-2260492-2022-22) and conducted in compliance with the Austrian Federal Act on Animal Welfare and the ethical standards of the responsible local committee and competent authorities. Sample sizes were determined based on prior experience with the model and were sufficient to detect biologically meaningful differences in tumor growth. Only female mice were used in accordance with the established orthotopic mammary fat pad model for hormone receptor-positive breast cancer.

Female athymic nude mice (Crl:NU(NCr)-Foxn1nu; Charles River), 8-12 weeks of age, were housed under pathogen-free conditions at 22 ± 1 °C, 55 ± 5% humidity and a 14 h light/10 h dark cycle. Autoclaved food, bedding and acidified drinking water were provided *ad libitum*.

To support estrogen-dependent tumor engraftment, mice were implanted subcutaneously with a 17β-estradiol slow-release pellet (0.36 mg, 90-day release; Innovative Research of America, #NE1210.36MG_90_50P) two days prior to tumor cell injection. Before injection, MCF7 iCas9 cells were transduced with lentivirus expressing 3xCTRL or 3xRNF sgRNAs alongside mCherry and puromycin resistance. Cas9 expression was induced with doxycycline (1 µg ml−1) and Cas9-BFP and sgRNA-mCherry double positive cells were isolated via FACS on a BD FACSAria Fusion. For orthotopic injections, sorted cells were expanded and resuspended in PBS mixed 1:1 with growth factor-reduced Matrigel (Corning) and 1 × 10⁷ cells were injected into the lower left mammary fat pad in a final injection volume of 100 µL. Tumors were monitored by caliper measurements every 3-7 days, and tumor volume was calculated according to the formula: volume = (D × d²)/2, in which D and d represent the long and short tumor diameters, respectively. The experimental unit was defined as an individual mouse.

When tumors reached a mean volume >200 mm³, mice were randomly assigned to treatment groups and treatment was initiated. Mice received either vehicle (castor oil containing 10% ethanol) or Fulvestrant (Faslodex; AstraZeneca) at 150 mg kg^-1^, administered subcutaneously twice during the first week and once weekly thereafter. Investigators were not blinded to treatment allocation during tumor measurements. No animals were excluded from analysis after randomization. Mice were monitored daily and euthanized when humane endpoints, for example, >20% body weight loss, signs of distress or pain, or volume ≥1500 mm³, were reached. At endpoint, tumors were excised and divided. One portion was fixed in 4% paraformaldehyde for histological analyses, the remaining tissue was kept on ice in PBS supplemented with 20 mM N-ethylmaleimide and processed for protein extraction and immunoblot analysis.

### Immunoblot

Tumors were manually sectioned on ice and 20-50 mg tissue was homogenized in 500 μL RIPA buffer supplemented with benzonase (1:1,000, Millipore, #70746), EDTA-free Halt protease inhibitor cocktail (1:100, Thermo Scientific, #11804111) and N-Ethylmaleimide (20 mM; Thermo, #11891335) in a TissueLyser II. Lysed samples were supplemented with 0.5% SDS and cleared by sonication on a Bioruptor Pico device (Diagenode) and centrifugation at maximum speed for 30 min at 4 °C. Protein concentrations were determined using the Pierce BCA protein assay kit (Thermo Fisher Scientific, PI23225) and equal amounts of protein were loaded for immunoblots.

### Immunohistochemistry

Formalin-fixed, paraffin-embedded tumor sections were deparaffinized, rehydrated and subjected to antigen retrieval in citrate buffer (pH 6.0). Endogenous peroxidase activity was quenched with 3% hydrogen peroxide. Sections were incubated with anti-Ki67 antibody (1:100, Abcam, ab16667), followed by HRP-conjugated polymer detection (DCS, #PD000POL-K) and DAB visualization (Abcam, ab64238). Slides were counterstained with hematoxylin, dehydrated, coverslipped and scanned using the Pannoramic 250 FLASH III Digital Scanner (3Dhistech) at 20x magnification. The viable tumor tissue was annotated by a veterinary pathologist blinded to group assignments and three separate areas per slide were individually analyzed for percentage of Ki67 positive nuclei using cell detection in QuPath software (v0.6.0) (*101*). For statistical analysis, the three areas per slide were averaged, data was batch-corrected to account for technical differences between staining runs and analyzed using a pooled two-way linear model with post-hoc pairwise comparisons in R Studio.

### AI and large language models

R code for statistical analyses and plotting of data was written with help of GPT-4o (OpenAI), Claude Sonnet 4.6 (Anthropic) or Claude Opus 4.8 (Anthropic). Python code for analysis and quantification of crystal violet colony formation images was written by Claude Opus 4.6 (Anthropic). Literature research and assembly of references was supported by Undermind.

**Fig. S1.**
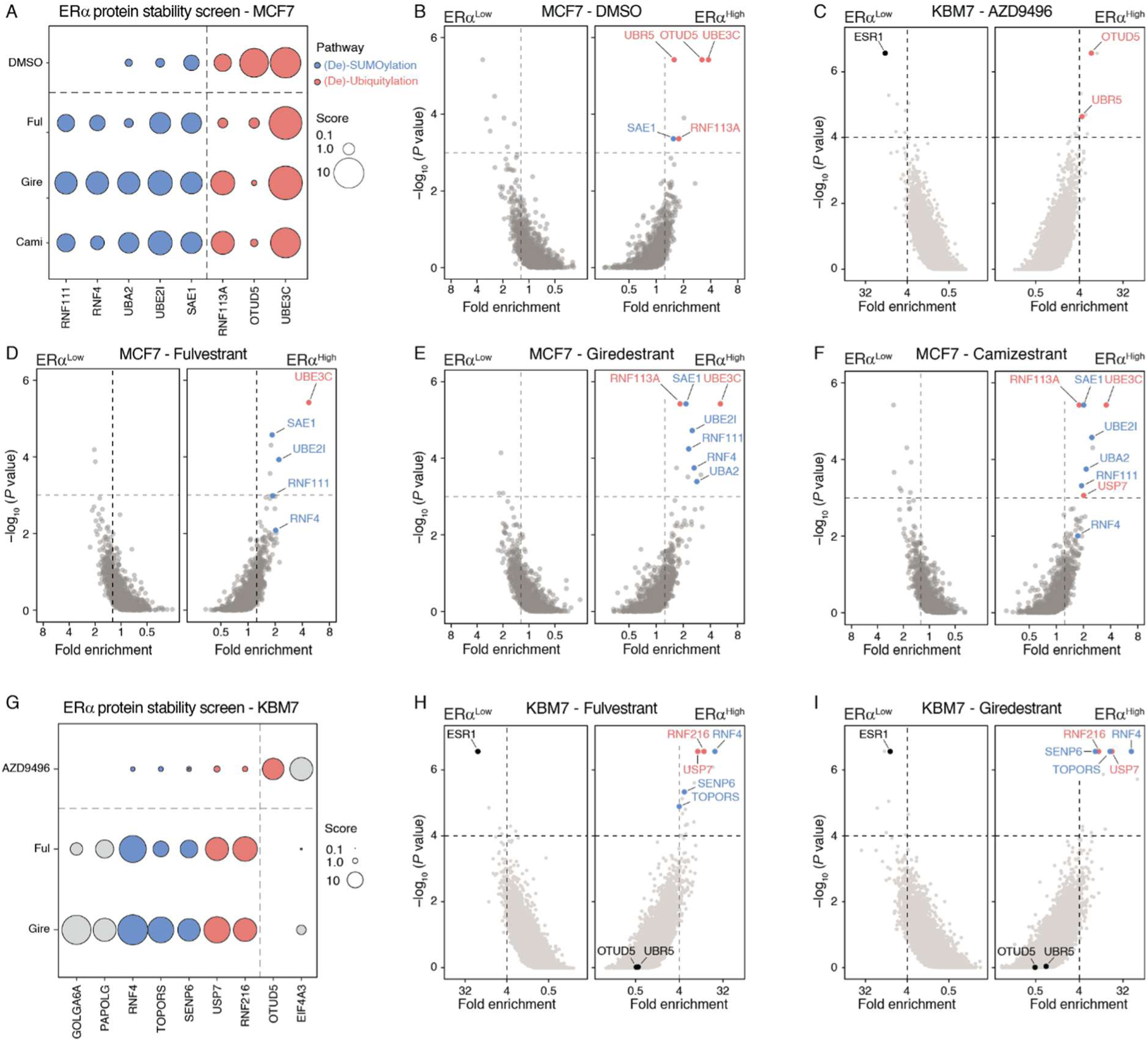
FACS-based CRISPR screens identify effectors of ERα degradation. **(A)** Enrichment scores of top screen hits for individual SERD and DMSO screens in MCF7 cells. Enrichment score was calculated as -Log_10_ (P value) × Log_2_ (fold-change). Hits with score > 4 in DMSO screen or combined analysis of Fulvestrant, Giredestrant and Camizestrant screens (Fig. 1C) are shown. Hits associated with SUMOylation or de-SUMOylation are shown in blue, hits associated with ubiquitylation or de-ubiquitylation in red. **(B)** FACS-based ERα protein stability CRISPR screen in MCF7 cells. MCF7 iCas9 ERα-GFP reporter cells treated with DMSO for 6 h were sorted based on ERα-GFP level normalized by mCherry, as in Fig. 1C. Combined MAGeCK (*84*) analysis of screens performed in n = 3 independent MCF7 iCas9 clones, gene-level fold enrichment (x axis) and one-sided MAGeCK P value (y axis). **(C)** Genome-wide ERα degradation CRISPR screen in KBM7 cells. KBM7 iCas9 cells expressing an ERα-BFP-P2A-mCherry stability reporter were mutagenized with a genome-wide sgRNA library (*77*) and treated with AZD9496 (500 nM) for 16 h before FACS isolation of ERα-BFP high and low fractions. MAGeCK analysis with n=2 technical replicates. **(D-F)** MCF7 ERα degradation CRISPR screens. MCF7 ERα reporter cells were mutagenized as in fig. S1B, treated with 10 nM Fulvestrant (D), Giredestrant (E) or Camizestrant (F) for 6 h and sorted based on ERα levels. MAGeCK analyses with n=3 technical replicates. **(G)** Enrichment scores of top KBM7 CRISPR screen hits. Enrichment score was calculated as in fig. S1A. Hits with enrichment score > 18 in AZD9496 screen (C) or combined analysis of Fulvestrant and Giredestrant screens (H and I, respectively) are shown. **(H-I)**, KBM7 genome-wide ERα degradation CRISPR screens. KBM7 ERα reporter cells were mutagenized as in fig. S1C, treated with 100 nM Fulvestrant (H) or Giredestrant (I) for 16 h and sorted based on ERα levels. MAGeCK analyses with n = 2 technical replicates as in fig. S1C.

**Fig. S2.**
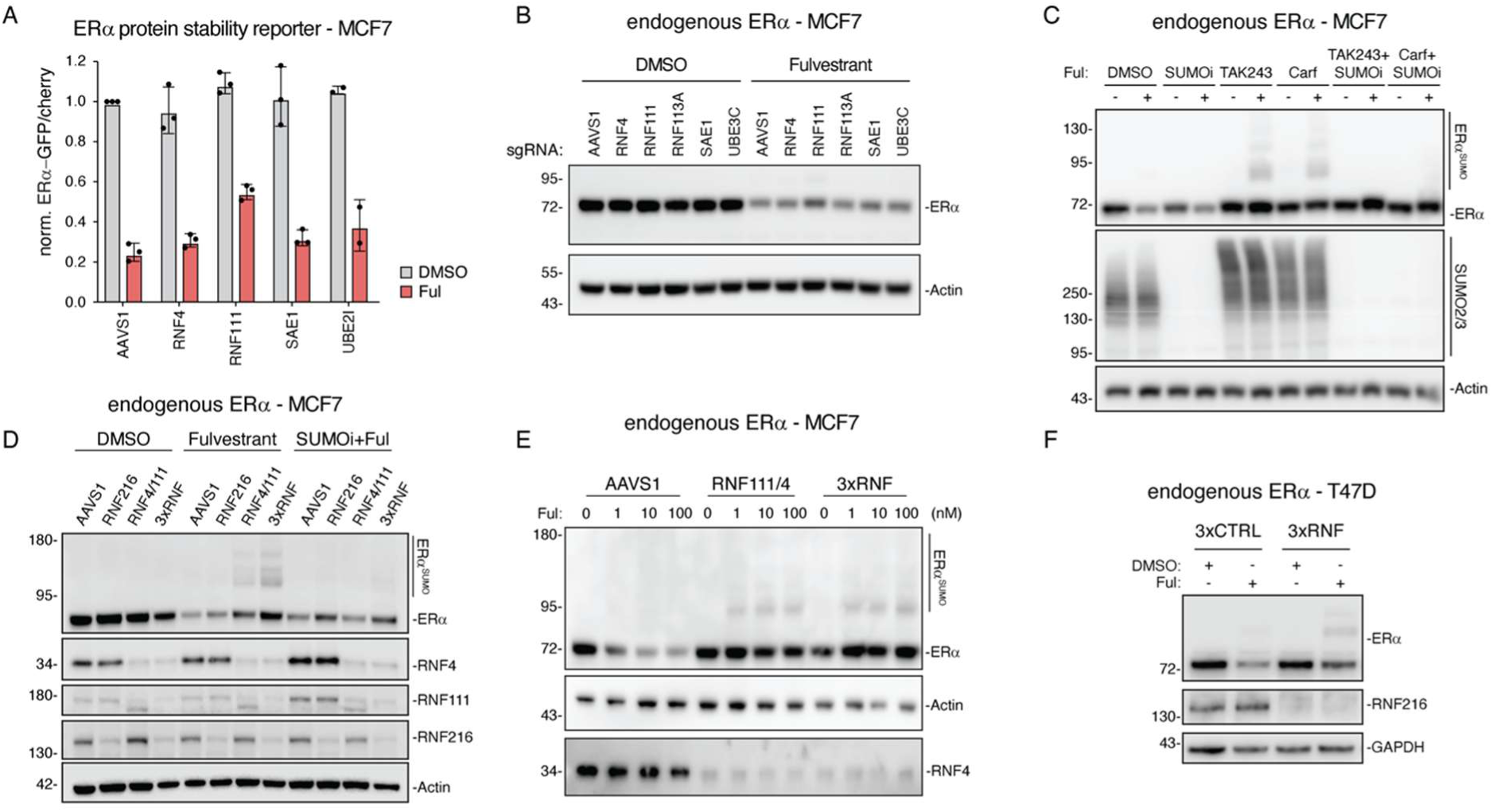
SERDs induce ERα degradation via SUMO-targeted ubiquitin E3 ligases. **(A)** FACS-based CRISPR screen validation. MCF7 iCas9 ERα-GFP reporter cells expressing the indicated sgRNAs were treated with DMSO or SERDs (10 nM) for 6 h before flow cytometric quantification. Mean ± sd. of n = 2 (UBE2I) or 3 (all others) independent experiments. **(B)** Immunoblot-based screen validation. MCF7 iCas9 cells expressing the indicated sgRNAs were treated with DMSO or Fulvestrant (10 nM) for 6 h before protein levels were analysed via immunoblot. Data representative of n = 2 independent experiments. **(C)** Immunoblot-based evaluation of SUMOylation. MCF7 iCas9 cells were pre-treated with DMSO, ML792 (SUMOi; 2 μM), TAK243 (ubiquitin activating enzyme inhibitor; 0.5 μM) or Carfilzomib (proteasome inhibitor; Carf; 1 μM) for 1 h, either alone or in combination, followed by treatment with or without Fulvestrant (10 nM) for 6 h. Data representative of n = 2 independent experiments. **(D)** RNF216-based degradation. MCF7 iCas9 cells were transduced with AAVS1-, RNF216-, combined RNF4- and RNF111- or combined RNF4-, RNF111- and RNF216-targeting (3xRNF) sgRNAs. Cells were pre-treated for 1 h with or without SUMOi (2 μM) followed by treatment with DMSO or Fulvestrant (10 nM) for 6 h as in Fig. 1F and ERα degradation was analyzed via immunoblotting. Data representative of n=3 independent experiments. **(E)** Combinatorial STUbL knockout. MCF7 iCas9 cells expressing AAVS1 CTRL, RNF4 and RNF111, or RNF4, RNF111 and RNF216 (3xRNF)-targeting sgRNAs were treated with indicated concentrations of Fulvestrant for 6 h before immunoblotting. Data representative of n = 2 independent experiments. **(F)** STUbL knockdown in T47D cells. T47D cells transduced with shRNA-miR vectors expressing non-targeting control (Renilla) or shRNAs targeting RNF4, RNF111 and RNF216 (3xRNF) were treated with or without Fulvestrant (10 nM) for 6 h before immunoblot analysis. Data representative of n=2 independent experiments. Ful, Fulvestrant; Cami, Camizestrant; Carf, Carfilzomib; 4-OHT, 4-Hydroxytamoxifen; STUbL, SUMO-targeted ubiquitin ligase; SUMOi, ML792.

**Fig. S3.**
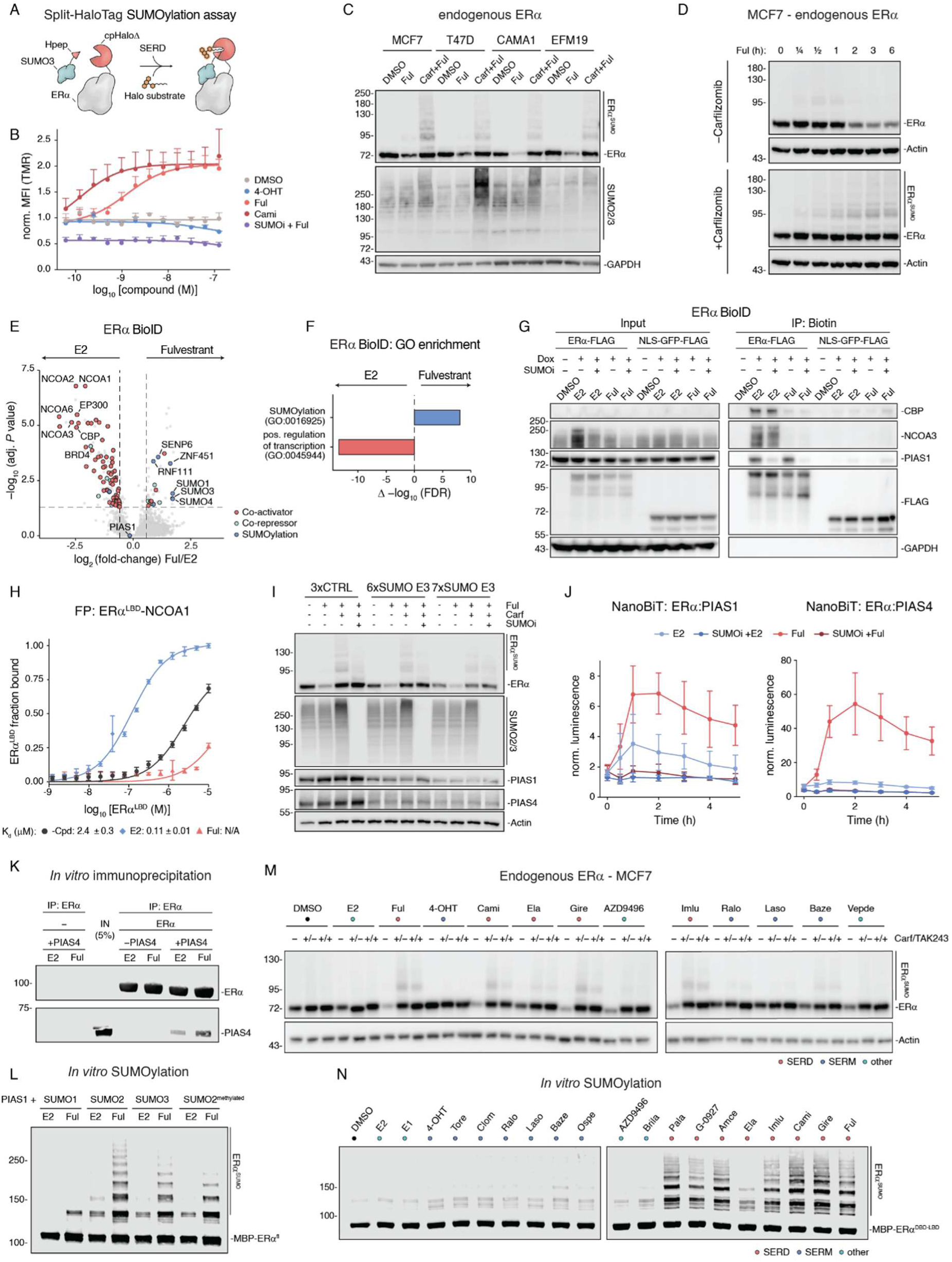
SERDs trigger ERα SUMOylation by the PIAS SUMO E3 ligase family. **(A**) Split-HaloTag assay schematic. Complementation of cpHaloΔ and Hpep peptide reconstitutes a functional HaloTag that covalently self-labels in the presence of TAMRA-conjugated Halo substrate. Co-expression of cpHaloΔ and Hpep as fusions to ERα and SUMO3, respectively, split-HaloTag complementation reports ERα SUMOylation in live cells. **(B)** Split-HaloTag-based ERα SUMOylation assay. HEK293T cells were co-transduced with cpHaloΔ-ERα and Hpep-SUMO3, pre-treated with Carfilzomib (1 μM) for 1 h in presence or absence of ML792 (SUMOi; 2 μM) followed by treatment with indicated concentrations of 4-OHT, Camizestrant or Fulvestrant for 6 h in the presence of TAMRA-conjugated Halo ligand. TAMRA labelling intensity was quantified via flow cytometry. n = 3 independent experiments, mean ± SEM. **(C)** ERα SUMOylation in breast cancer cell lines. ERα^+^ breast cancer cell lines (MCF7, T47D, CAMA1, EFM19) were treated with DMSO or Fulvestrant (100 nM) for 6 h with or without 1 h Carfilzomib (1 μM) pre-treatment. Levels of ERα SUMOylation and total SUMOylated proteins were evaluated via immunoblotting. Data is representative of n = 3 independent experiments. **(D)** SUMOylation kinetics. MCF7 cells were pre-treated with or without Carfilzomib (1 μM, 1 h), followed by treatment with Fulvestrant (10 nM) for increasing amounts of time. Levels of ERα degradation (top) and ERα SUMOylation (bottom) were evaluated via immunoblotting. **(E-G)** ERα BioID proximity labelling. ERα-miniTurbo cells were treated with 10 nM Fulvestrant or Estradiol as in Fig. 2C and D and ERα interactors quantified via quantitative proteomics (E and F) or immunoblotting (G). **(E)** Transcriptional activators (red; GOBP:0045944) and SUMO pathway components (blue; GOBP:0016925) in the scoring window (FC > 1.5, adj. P value < 0.05, one-way ANOVA with Tukey’s post hoc test and Benjamini-Hochberg correction, n = 3 biological replicates) are highlighted. **(F)** GO analysis of ERα BioID interactors. Top GO biological process terms overrepresented among ERα interactors upon Fulvestrant versus Estradiol treatment (Δ -log_10_(FDR); Benjamini-Hochberg-corrected two-tailed Fisher’s exact test). **(G)** ERα- and NLS-GFP-miniTurbo cells were treated with Fulvestrant or Estradiol (10 nM each) in the presence or absence of SUMOi ML792 (2 µM) and selected interactors were visualized via immunoblotting. Data representative of n = 3 independent experiments. **(H)** Fluorescence polarization assay. Binding of ERα^LBD^ to a fluorescein-labelled NCOA1 peptide in the presence of Estradiol, Fulvestrant or no ligand. Values are shown as fraction bound, mean +/- sd. of n = 3 technical replicates. **(I)** Cellular ERα SUMOylation. MCF7 iCas9 cells expressing 3xCTRL sgRNAs, a combination of sgRNAs targeting PIAS1-4 and ZMIZ1-2 (6xSUMO E3) or PIAS1-4, ZMIZ1-2 as well as ZNF451 (7xE3). Data representative of n = 3 independent experiments. **(J)** NanoBiT ERα interaction assay. HEK293 cells were co-transfected with ERα-LgBiT and the indicated SmBiT fusion (PIAS1, PIAS4), pre-treated with Carfilzomib (1 µM) with or without SUMOi ML792 (2 µM) in the presence of Endurazine live cell substrate, followed by treatment with Estradiol or Fulvestrant (10 nM each). Bioluminescence reporting split-NanoLuc reconstitution was recorded over 5 h post-treatment. n = 2 independent experiments, mean ± SD. **(K)** *In vitro* ERα pulldowns. STREP-MBP-ERα^fl^ was immobilized on STREP beads and subsequently incubated with or without PIAS4^fl^ in the presence of E2 or Fulvestrant, followed by immunoblotting. Data representative of n = 3 independent experiments. **(L)** *In vitro* ERα SUMOylation. MBP-ERα^fl^ was incubated with PIAS1 and indicated SUMO isoform or methylated SUMO2 incapable for forming poly-SUMO chains, in the presence of Estradiol or Fulvestrant, followed by visualization via immunoblotting. Data representative of n = 2 independent experiments. **(M)** Cellular ERα SUMOylation. MCF7 WT cells were treated with the indicated ERα ligands (10 nM each) for 6 h in with or without Carfilzomib (1 µM) and/or SUMOi (2 µM) pre-treatment. Levels of SUMOylated and non-SUMOylated ERα were quantified via immunoblotting. Data representative of n = 2 independent experiments. **(N)** *In vitro* ERα SUMOylation. MBP-ERα^DBD-LBD^ was incubated with SUMO2/3, PIAS4 and indicated ERα ligands (10 µM each). SUMOylated ERα was visualized via ERα immunoblotting. Data representative of n = 2 independent experiments. Quantification of (M) and (N), Fig. 2F. MBP, Maltose-binding protein; -Cpd, without compound; fl, ERα full-length protein; LBD, isolated ERα ligand binding domain; DBD-LBD, ERα DNA binding domain fused to ligand binding domain; Amce, Amcenestrant; Baze, Bazedoxifene; Brila, Brilanestrant; Cami, Camizestrant; Clomi, Clomiphene; Ela, Elacestrant; E1, Estrone; E2, Estradiol; Ful, Fulvestrant; G-0927, GDC-0927; Gire, Giredestrant; Imlu, Imlunestrant; Laso, Lasofoxifene; Ospe, Ospemifene; Pala, Palazestrant; Ralo, Raloxifene; Tore, Toremifene; 4-OHT, 4-Hydroxytamoxifen; Carf, Carfilzomib.

**Fig. S4.**
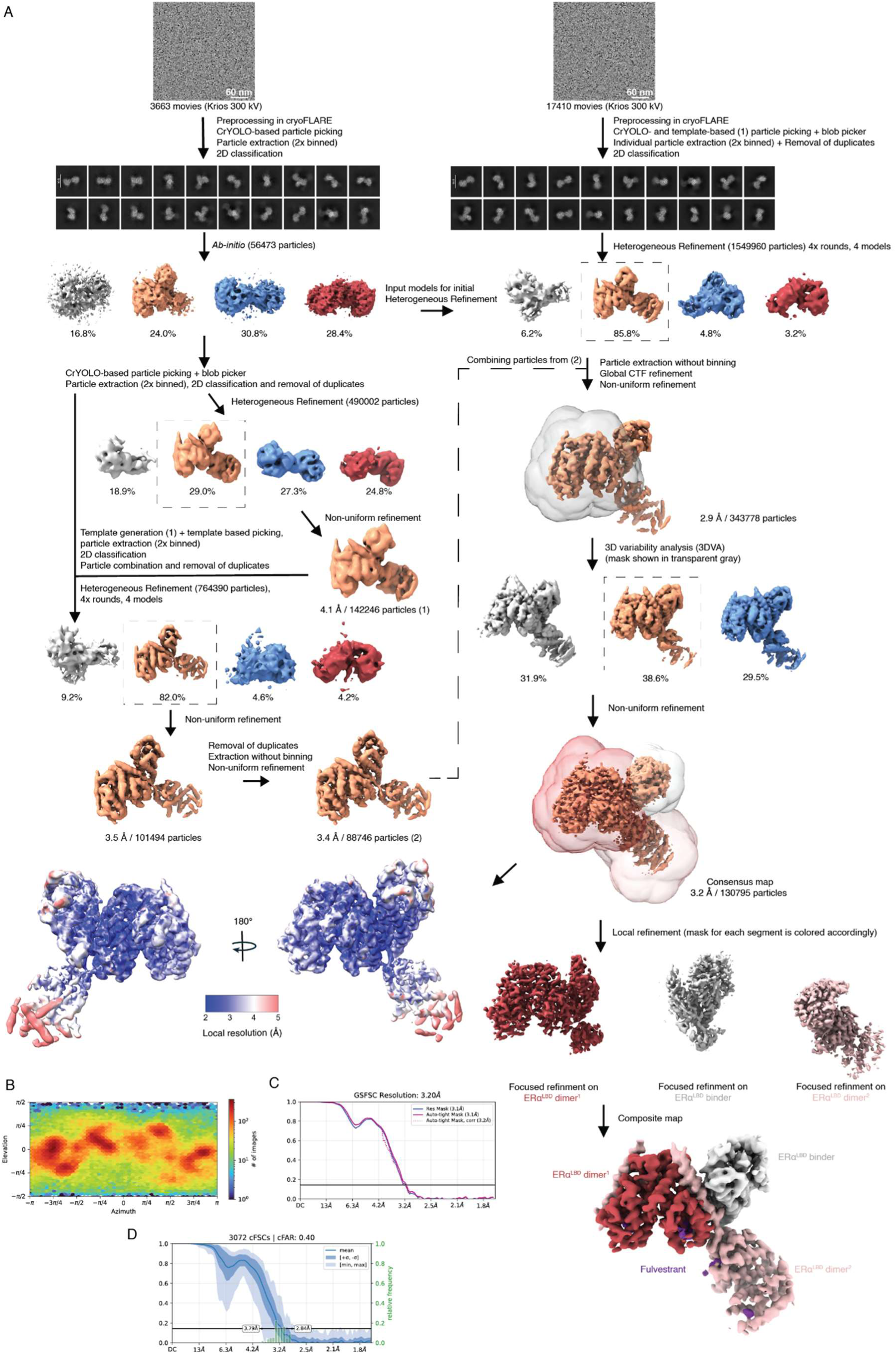
Cryo-EM data processing of tetrameric BRIL-ERαLBD:Fulvestrant:de novo ERαLBD binder complex. **(A**) Summary of cryo-EM data processing pipeline. The consensus map is presented in two orientations and colored based on local resolution. **(B)** Angular distribution plot of particles used to generate the consensus map. **(C)** Gold-standard Fourier shell correlation (GSFSC) curves after FSC mask auto-tightening at the threshold of 0.143. **(D)** Conical FSC area ratio (cFAR) for the consensus map.

**Fig. S5.**
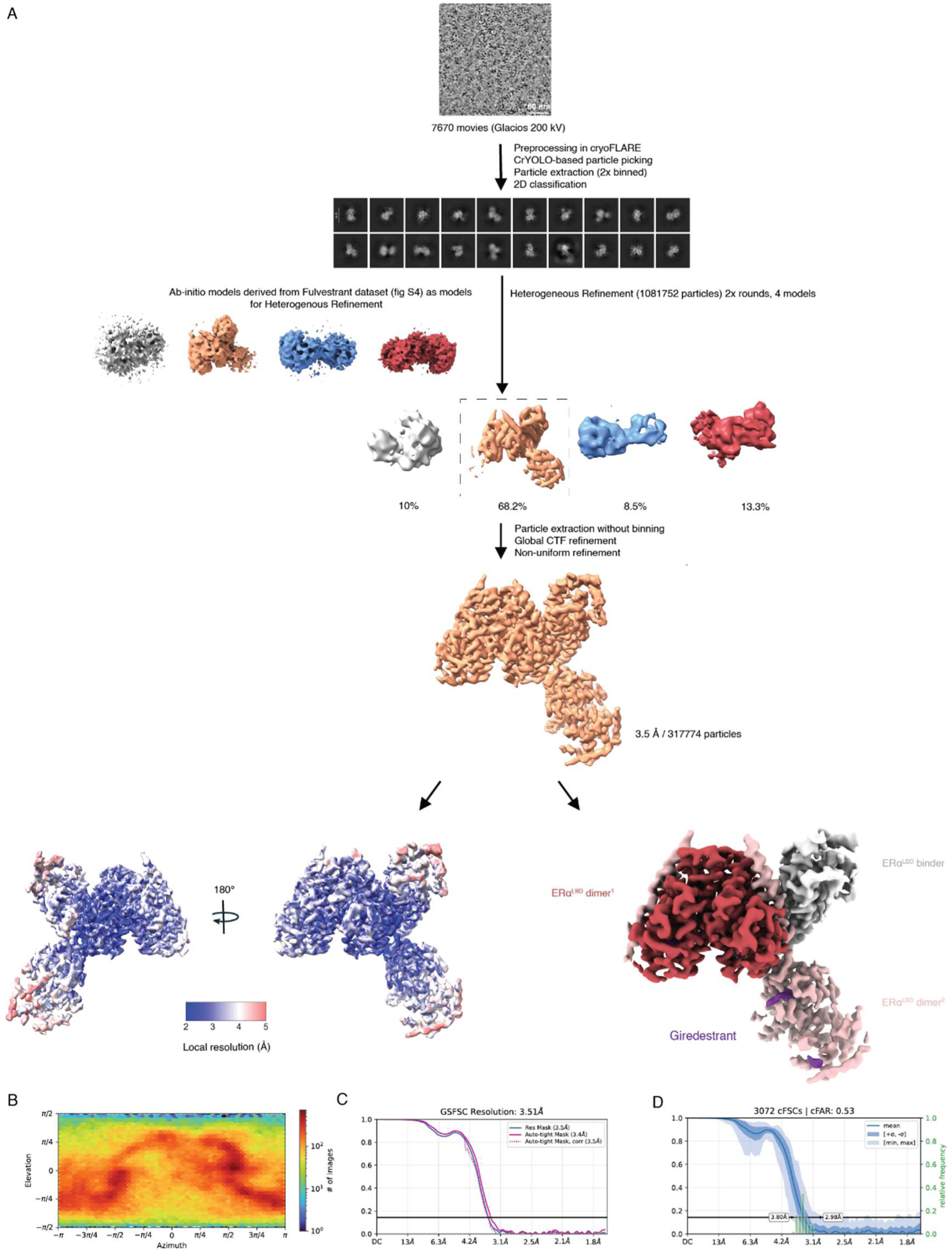
Cryo-EM data processing of tetrameric BRIL-ERαLBD:Giredestrant:de novo ERαLBD binder complex. **(A)** Summary of the Cryo-EM data processing pipeline. The final map after non-uniform refinement is presented in two orientations and colored by local resolution. **(B)** Angular distribution of particles used for final non-uniform refinement. **(C)** Gold-standard Fourier shell correlation (GSFSC) curves after FSC mask auto-tightened at the threshold of 0.143. **(D)** Conical FSC area ratio (cFAR) for final density map.

**Fig. S6.**
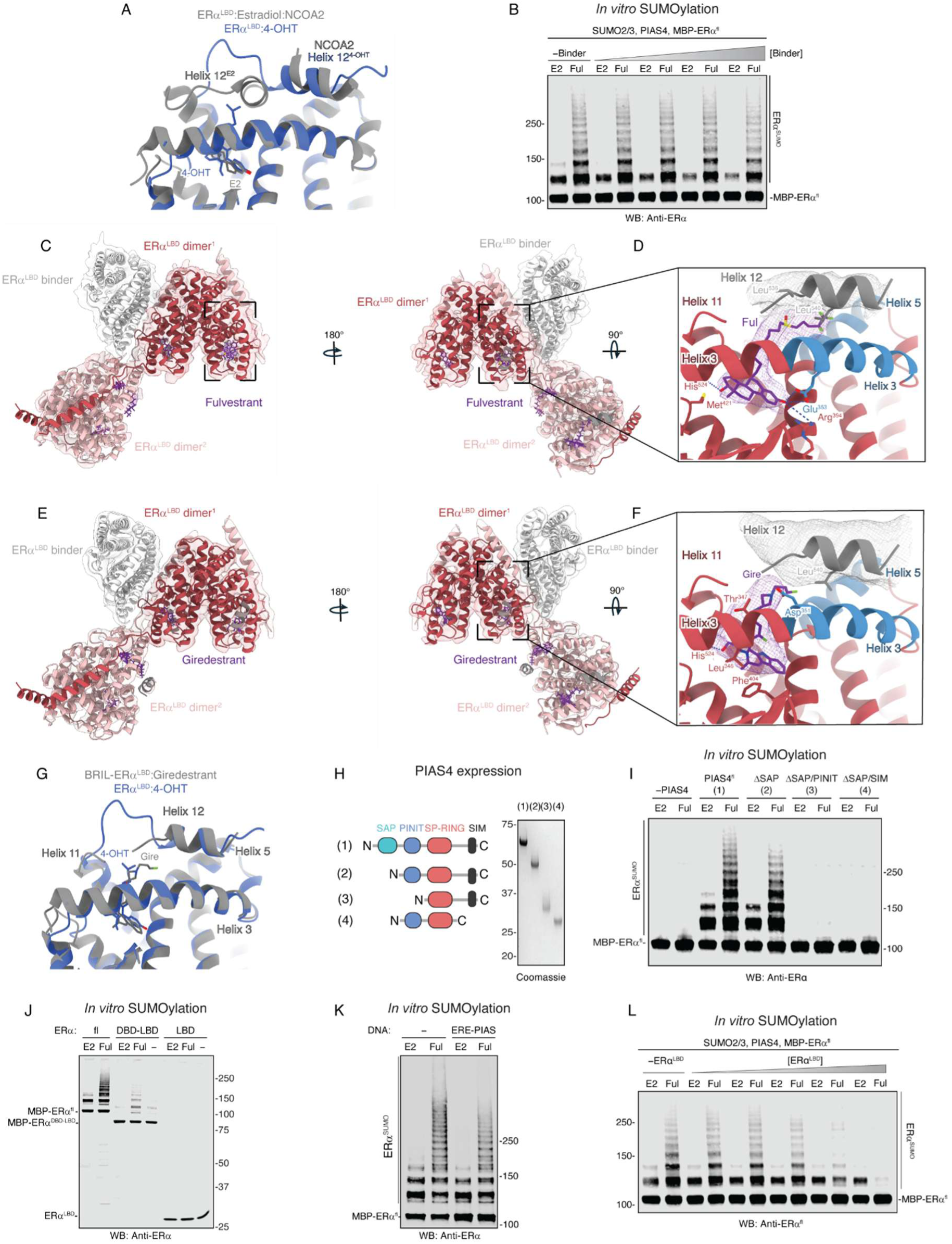
SERD-SUMOylation requires the ERα DNA binding domain. **(A)** Comparison of agonist and antagonist ligand-binding modes. ERα^LBD^ bound to Estradiol (E2) and an NCOA2 peptide (PDB: 1GWR, gray) superimposed with ERα^LBD^ bound to 4-Hydroxytamoxifen (4-OHT) (PDB: 3ERT, blue). Different helix 12 (H12) conformations are highlighted. **(B)** *In vitro* ERα SUMOylation. Full-length MBP-ERα (MBP-ERα^fl^) was incubated with SUMO2/3 and PIAS4 in the presence of Estradiol or Fulvestrant and increasing concentrations of the *de novo* designed ERα^LBD^ binder used for cryo-EM studies. SUMOylated ERα was visualized via ERα immunoblotting. Data representative of n = 2 independent experiments. **(C-F)** Cryo-EM of tetrameric BRIL-ERα^LBD^ bound to Fulvestrant or Giredestrant with a de novo ERα^LBD^ binder. BRIL-ERα^LBD^ oligomers form via Helix 1-mediated domain-swapping between ER dimers, regardless of the binder (data not shown). High-resolution structures were obtained only in the presence of binder, with two LBD dimers and one binder constituting the repeating unit. **(C)** Composite cryo-EM map of the full tetrameric BRIL-ERα^LBD^ (red) complex bound to Fulvestrant (purple) and a *de novo* ERα^LBD^ binder (gray), overlaid with the corresponding atomic model and the outlined ligand-binding site shown in Fig. 2G-I. **(D)** Close-up view of the ligand-binding site with residual helix 12 density (gray) outlined in (C), highlighting a steric clash between Leu540 in H12 and the Fulvestrant (Ful) tail upon static modelling. Protein side chains interacting with Fulvestrant (d < 4 Å) are shown with hydrogen bonds represented by dotted lines. Cryo-EM density for Fulvestrant and H12 is shown as purple and gray mesh, respectively. The co-activator binding groove is colored in blue. **(E)** Cryo-EM map of the full tetrameric BRIL-ERα^LBD^ (red) complex bound to Giredestrant (purple) and a *de novo* ERα^LBD^ binder (gray), overlaid with the corresponding atomic model. **(F)** Close-up view of one ERα^LBD^ monomer outlined in (E). In the presence of Giredestrant (purple), H12 (gray) assumes a canonical antagonist-bound conformation across all compound binding site. Protein side chains interacting with Giredestrant (d < 4 Å) are shown with hydrogen bonds represented by dotted lines. Cryo-EM density for Giredestrant and H12 is shown as purple and gray mesh, respectively. The co-activator binding groove is colored in blue. **(G)** Overlay of the Giredestrant binding mode (gray) shown in (F) superimposed with the X-ray structure of ERα^LBD^ bound to 4-OHT (PDB: 3ERT, blue). The residues 341-545 of both structures were aligned to determine an RMSD of 1.5 Å over 198 residues. **(H)** Recombinant expression of PIAS4 mutants. Full length PIAS4 (PIAS4^fl^; 1) or variants harboring deletions of the SAP domain (2), the SAP and PINIT domains (3) or the SAP domain and C-terminal SIM (4) were recombinantly expressed and purified for *in vitro* SUMOylation assays. **(I-L)** *In vitro* ERα SUMOylation assays. **(I)** MBP-ERα^fl^ was incubated with SUMO2/3 and with or without PIAS4 variants in the presence of Estradiol or Fulvestrant. **(J)** MBP-ERα^fl^ or ERα truncations consisting of the DNA- and ligand-binding domain (DBD-LBD) or only the LBD were incubated with PIAS4 and SUMO2/3 as well as Estradiol, Fulvestrant or no ligand. **(K)** MBP-ERα^fl^ was incubated with SUMO2/3, PIAS4 and Estradiol or Fulvestrant in the presence or absence of a DNA oligonucleotide containing an estrogen response element fused to a PIAS-binding site (*87*). **(L)** MBP-ERα^fl^ was incubated with SUMO2/3, PIAS4 and increasing concentrations of LBD in the presence of Estradiol or Fulvestrant. Data representative of n = 3 (I) or n = 2 (J-L) independent experiments with SUMOylated ERα visualized via immunoblotting ERE, Estrogen response element.

**Fig. S7.**
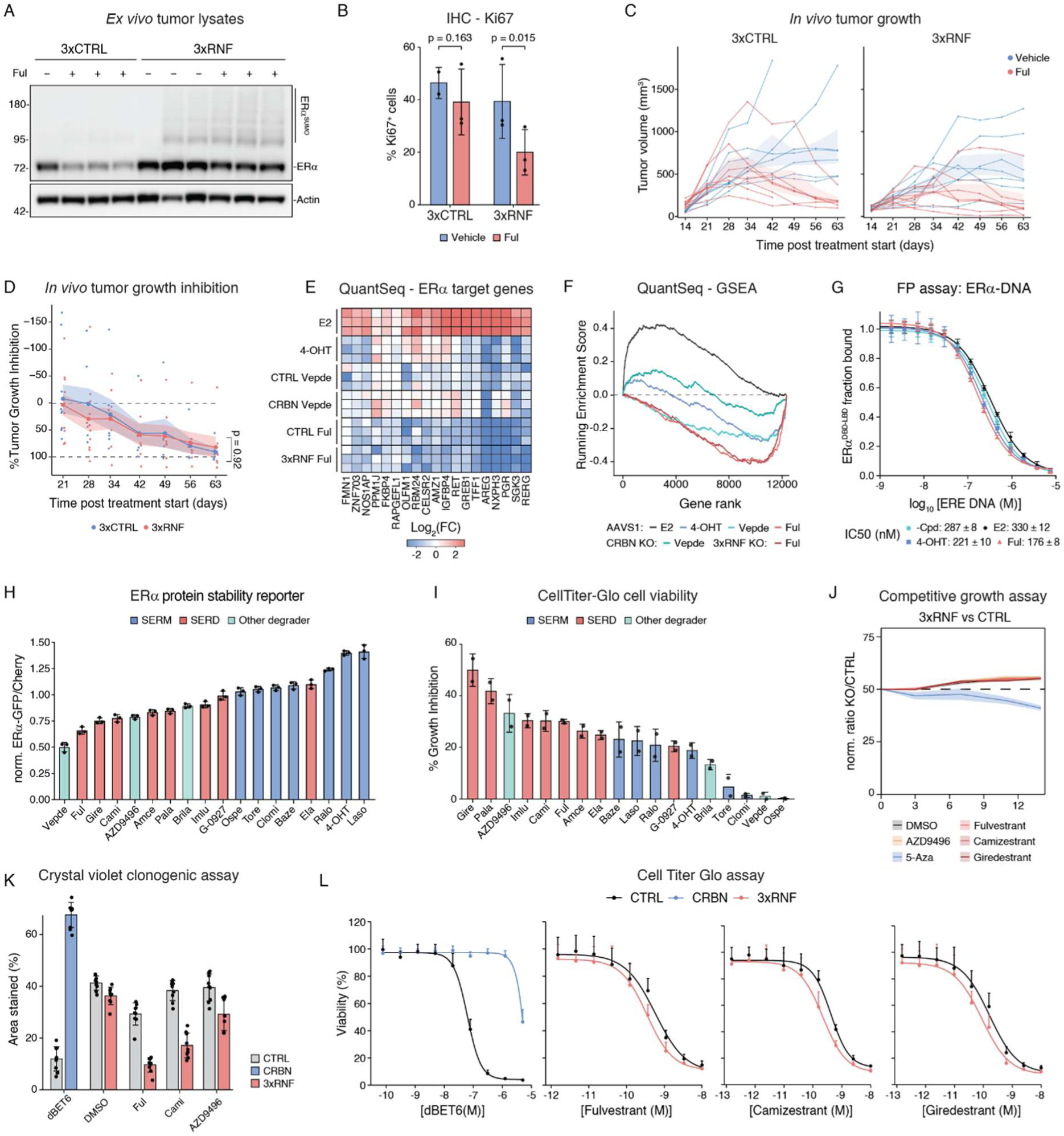
ERα degradation is not required for SERD activity. **(A)** Immunoblot-based quantification of ERα stability in orthotopic xenografts. MCF7 iCas9 3xCTRL or 3xRNF KO cells were injected into athymic nude mice and after tumor establishment, mice were treated with two doses of vehicle or Fulvestrant (150 mg/kg) before tumors were harvested for immunoblotting. Samples represent tumors from different mice. Data representative of n = 2 independent experiments. **(B)** Ki67 expression. Xenografts were established and mice treated as in (A). Tumors were immunohistochemically stained for Ki67 expression. Mean ± sd. of n = 2 (3xCTRL + Vehicle) or 3 (all others) tumors, pooled two-way linear model with post-hoc pairwise comparisons. **(C)** *In vivo* tumor growth. MCF7 iCas9 3xCTRL or 3xRNF KO orthotopic xenograft models in athymic nude mice were established as in (A) and vehicle or Fulvestrant (150 mg/kg) were administered subcutaneously once per week. Tumor size was measured at indicated timepoints after cell injection. Lines, growth curves of individual tumors from Fig. 3B; n at start of treatment = 8 (CTRL vehicle), 10 (CTRL Fulvestrant), 7 (3xRNF vehicle), 10 (3xRNF Fulvestrant) individual mice; ribbons, SEM. **(D)** *In vivo* tumor growth inhibition. Per-mouse percent growth inhibition by Fulvestrant compared to vehicle was calculated for 3xCTRL (blue) and 3xRNF tumors (red). Dots represent individual tumors from (C). Lines and ribbons, mean +/- SEM. Two-way ANOVA on log-transformed AUC (tumor volume; Fig. 3B). **(E-F)** Transcriptional profiling of ERα activity. MCF7 iCas9 CTRL, CRBN or 3xRNF knockout cells were treated with DMSO or Estradiol, 4-OHT, Fulvestrant (all 10 nM) or Vepdegestrant (100 nM) for 24 h in n = 3 biological replicates before transcriptional profiling *via* QuantSeq. Estradiol-treated cells were deprived of hormones for 2 days prior to treatment. Log_2_ fold-changes of core estrogen target genes (*11*) (**E**) and gene set enrichment analysis of MSigDB Hallmark ESTROGEN_RESPONSE_EARLY (*50*) genes (**F**). **(G)** Fluorescence polarization assay. Competitive reverse titration with unlabeled ERE oligonucleotide against a preformed complex of MBP-ER^DBD-LBD^ with the same fluorescein-labelled ERE oligonucleotide in the presence of Estradiol, Fulvestrant or no ligand. Values are shown as fraction bound, mean +/- sd. of n = 3 technical replicates. **(H)** ERα degradation potency. MCF7 iCas9 ERα-GFP reporter cells were treated with the indicated compounds (all 10 nM) for 6 h, and ERα-GFP signal normalized by iRFP-H2A was quantified by FACS. Mean ± sd. of n = 3 independent experiments. **(I)** Percent growth inhibition. MCF7 iCas9 cells were treated with increasing doses of indicated compounds for 7 days and cell viability was evaluated via CellTiterGlo assay. Mean ± sd. of n = 2 independent experiments with 3 biological replicates each. **(J)** Competitive growth assay in RKO cells. GFP expressing RKO iCas9 3xCTRL cells were mixed with mCherry expressing 3xRNF knockout cells and treated with DMSO, AZD9496 (0.5 nM), Fulvestrant (0.5 nM), Camizestrant (0.25 nM), Giredestrant (0.25 nM) or 5-Azacytidine (1 μM). Ratio of mCherry^+^/GFP^+^ cells was evaluated in regular intervals via flow cytometry. Lines and ribbons, mean ± SEM for n = 2 (5-Azacytidine) or 3 (all others) independent experiments. **(K)** Colony formation assay. MCF7 iCas9 control, CRBN KO, or 3xRNF KO cells were treated with DMSO, Fulvestrant (0.5 nM), Camizestrant (0.25 nM), AZD9496 (0.5 nM), or dBET6 (250 nM) for 13 days before crystal violet staining. Mean ± sd. of n = 3 independent experiments with 3 biological replicates each. Representative images are shown in Fig. 3D. **(L)** Cell growth assay. MCF7 iCas9 control, CRBN or 3xRNF knockout cells were treated with increasing doses of dBET6, THAL-SNS-032, Fulvestrant or Camizestrant for 7 days and cell viability was evaluated via CellTiterGlo assay. Mean ± sd. of n = 2 (dBET6, THAL-SNS-032) or 3 (Fulvestrant, Camizestrant) independent experiments with 3 biological replicates each. In (K) and (L) CRBN KO was compared to AAVS1; 3xRNF KO was compared to 3xCTRL. ERE, Estrogen response element; -Cpd, without compound; MBP, Maltose-binding protein; Amce, Amcenestrant; Baze, Bazedoxifene; Brila, Brilanestrant; Cami, Camizestrant; Clomi, Clomiphene; Ela, Elacestrant; E2, Estradiol; Ful, Fulvestrant; G-0927, GDC-0927; Gire, Giredestrant; Imlu, Imlunestrant; Laso, Lasofoxifene; Ospe, Ospemifene; Pala, Palazestrant; Ralo, Raloxifene; Tore, Toremifene; 4-OHT, 4-Hydroxytamoxifen.

**Fig. S8.**
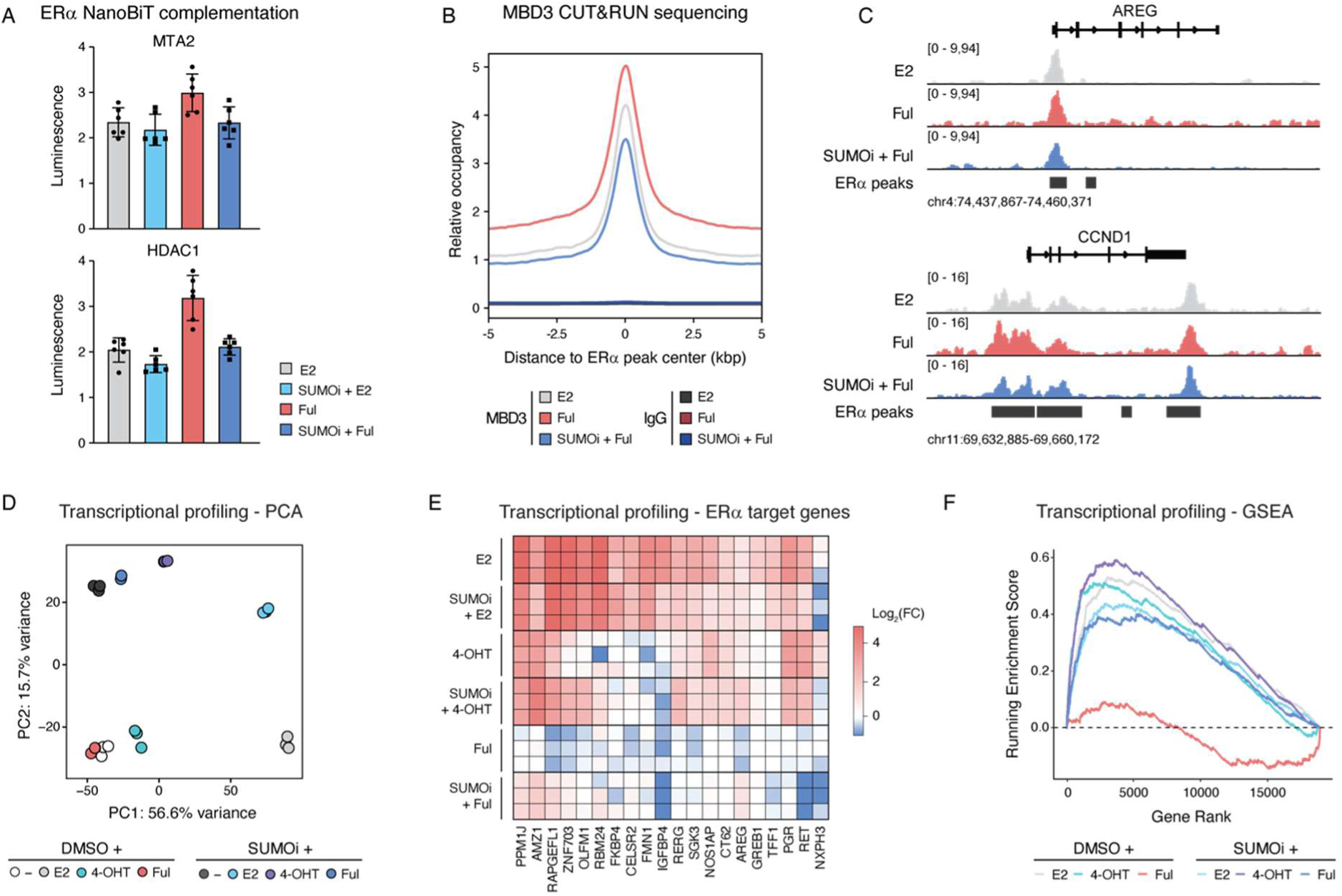
SUMOylation is required for full antiestrogenic effects of SERDs. **(A)** NanoBiT ERα interaction assay. HEK293 cells were co-transfected with ERα-LgBiT and the indicated SmBiT fusion (MTA2, HDAC1), pre-treated with Carfilzomib (1 µM) with or without SUMOi ML792 (2 µM) in the presence of Endurazine live cell substrate, followed by treatment with Estradiol or Fulvestrant (10 nM each). Bioluminescence reporting split-NanoLuc reconstitution was recorded after 2 h of treatment. n = 2 independent experiments, mean ± SD. **(B-C)** CUT&RUN sequencing of NuRD chromatin occupancy. Average CUT&RUN signal profiles upon Estradiol, Fulvestrant (10 nM each) and SUMOi (ML792, 2 µM) + Fulvestrant treatment centered on Fulvestrant-induced ERα peak centers (±5 kb, 50-bp bins), shown as the mean across all peaks per sample. **(B)** Genome browser tracks of MBD3 CUT&RUN signal at the *AREG* and *CCND1* loci. Called ERα binding sites upon Fulvestrant treatment (ERα peaks) are shown as black bars beneath the coverage tracks. **(C) (D-F)** Transcriptional profiling of ERα activity. After 72 h hormone deprivation MCF7 cells were treated with DMSO, Estradiol, 4-OHT, Fulvestrant (all 10 nM), with or without SUMOi (ML792, 2 µM) pre-treatment, for 24 h in n = 3 biological replicates before transcriptional profiling via RNA-seq. Principal component analysis (**D**) Log_2_ fold-changes of core estrogen target genes (*11*) (**E**) and gene set enrichment analysis of MSigDB Hallmark ESTROGEN_RESPONSE_EARLY(*50*) genes (**F**). H12, Helix12; E2, Estradiol; Ful, Fulvestrant, SUMOi, ML792; 4-OHT, 4-Hydroxytamoxifen.

**Table S1.**
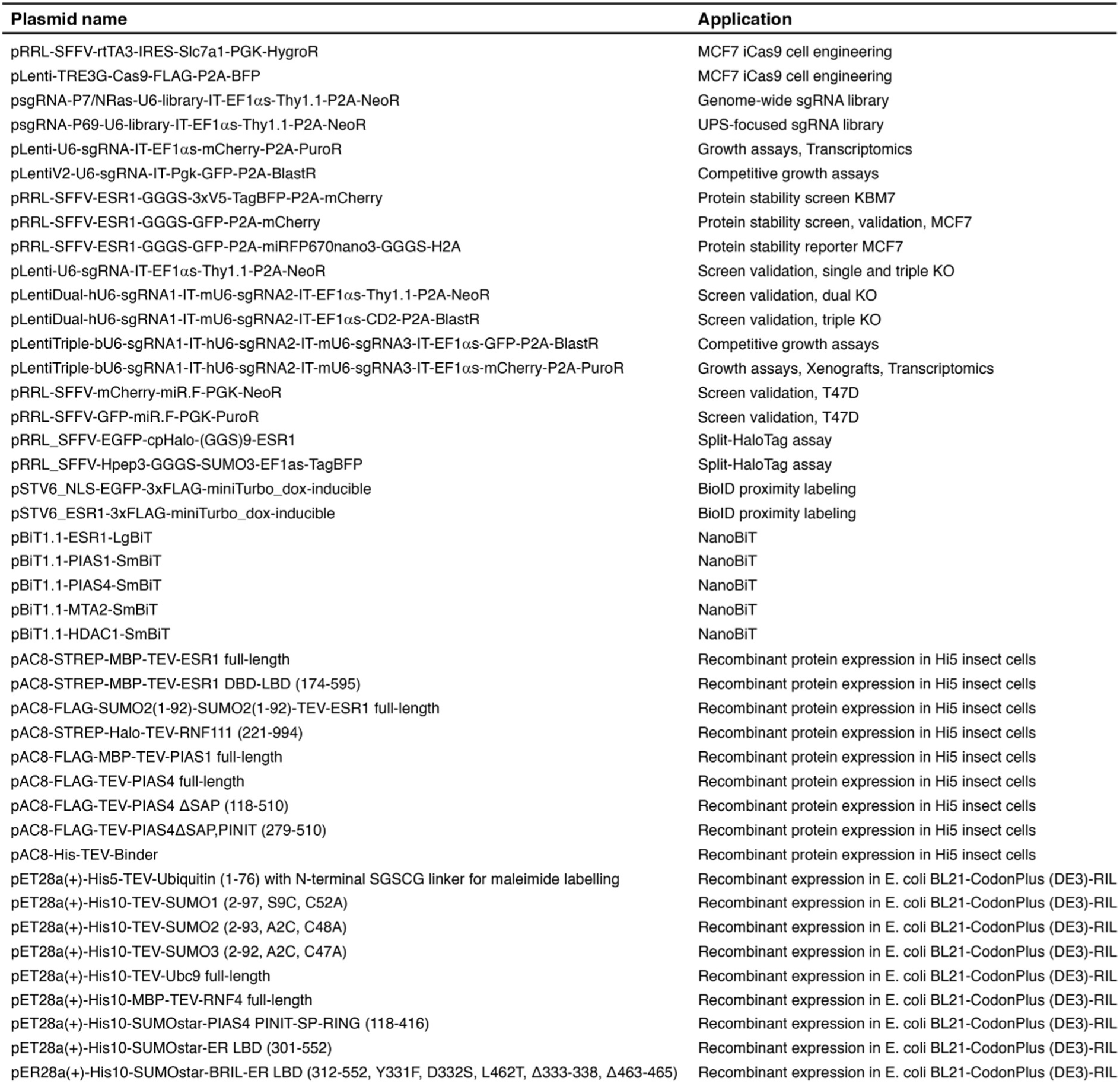
Plasmids used in this study.

**Table S2.**
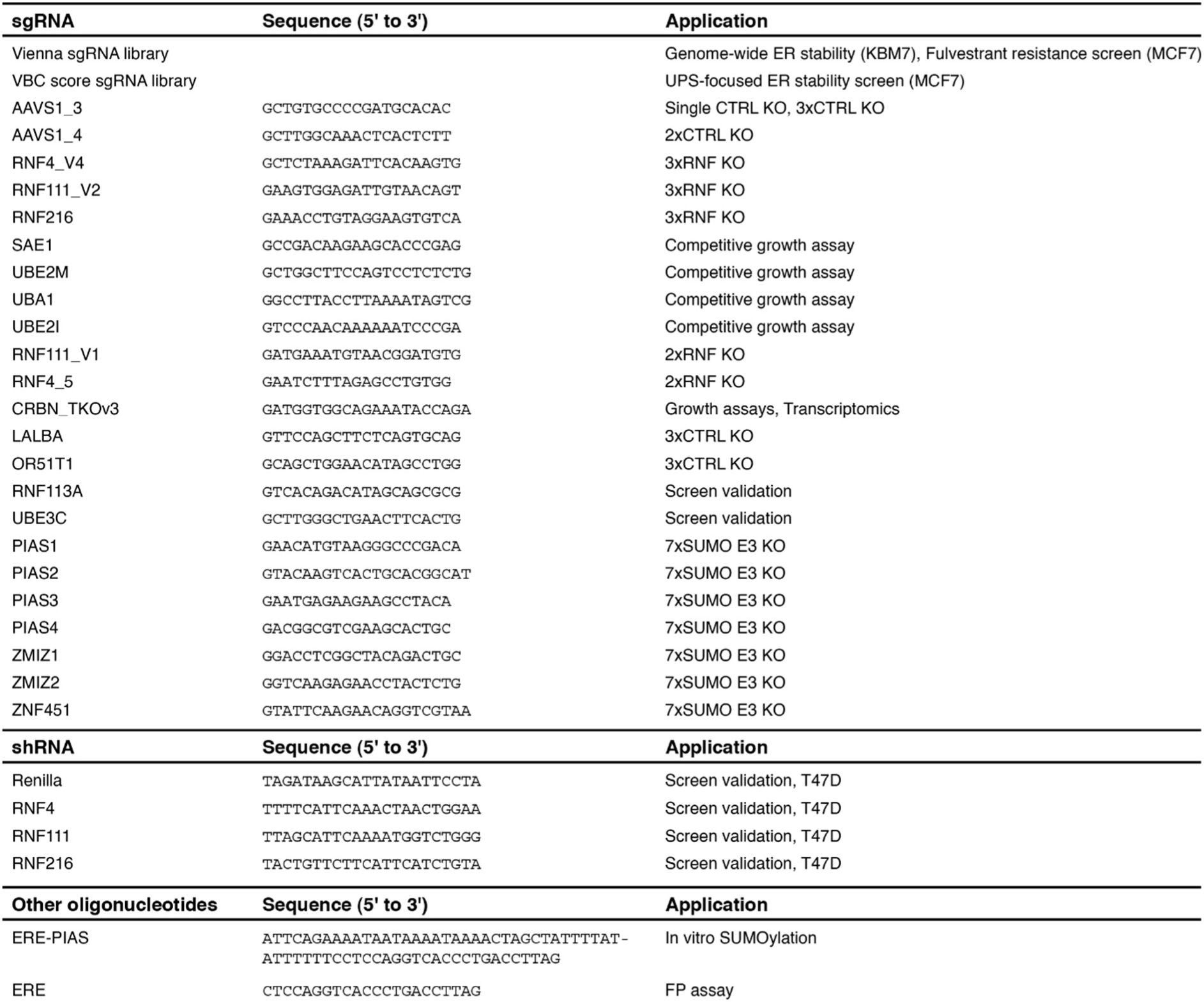
Oligonucleotides used in this study.

**Table S3.**
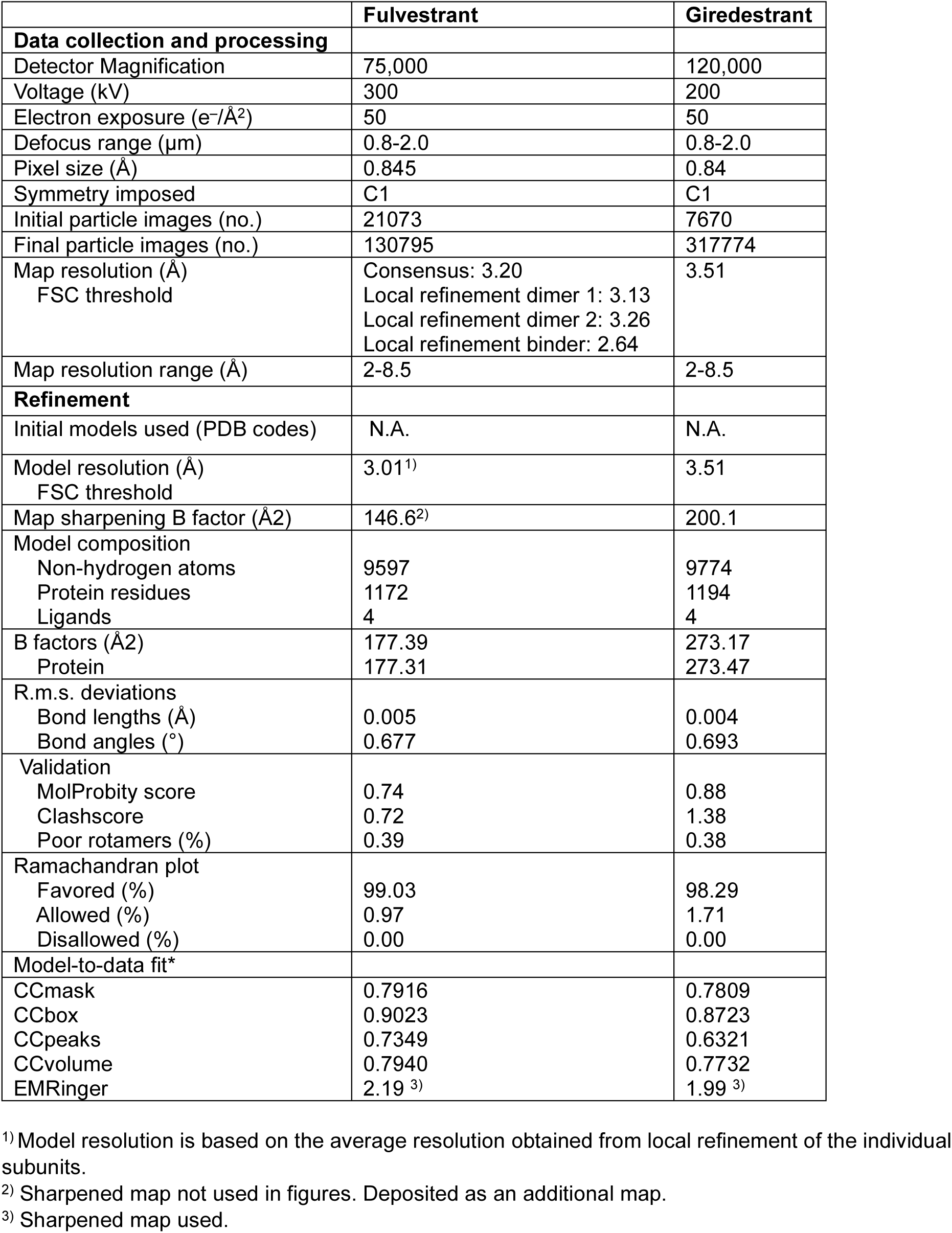
Cryo-EM data collection, refinement and validation statistics.

